# Single-cell transcriptomic analysis of antiviral responses and viral antagonism in Chikungunya virus-infected synovial fibroblasts

**DOI:** 10.1101/2020.06.07.138610

**Authors:** Fabian Pott, Dylan Postmus, Richard J. P. Brown, Emanuel Wyler, Elena Neumann, Markus Landthaler, Christine Goffinet

## Abstract

In recent years, (re-)emerging arboviruses including Chikungunya virus (CHIKV) and Mayaro virus (MAYV) have caused growing concern due to expansion of insect vector ranges. No protective vaccine or specific antiviral strategies are currently available. Long-term morbidity after CHIKV infection includes debilitating chronic joint pain, which has associated health and economic impact. Here, we analyzed the early cell-intrinsic response to CHIKV and MAYV infection in primary human synovial fibroblasts. This interferon-competent cell type represents a potential source of polyarthralgia induced by CHIKV infection. Synovial fibroblasts from healthy and osteoarthritic donors were similarly permissive to CHIKV and MAYV infection *ex vivo*. Using RNA-seq, we defined a CHIKV infection-induced transcriptional profile with several hundred interferon-stimulated and arthralgia-mediating genes upregulated. Type I interferon was both secreted by infected fibroblasts and protective when administered exogenously. IL-6 secretion, which mediates chronic synovitis, however, was not boosted by infection. Single-cell RNA-seq and flow cytometric analyses uncovered an inverse correlation of activation of innate immunity and productive infection at the level of individual cells. In summary, primary human synovial fibroblasts serve as bona-fide *ex vivo* primary cell model of CHIKV infection and provide a valuable platform for studies of joint tissue-associated aspects of CHIKV immunopathogenesis.

## Introduction

Chikungunya virus (CHIKV) and Mayaro virus (MAYV) are arthritogenic alphaviruses of the *Togaviridae* family, which are transmitted by *Aedes sp.* mosquitoes and circulate both in urban cycles between vectors and humans, and in sylvatic cycles [1–3]. Beyond the typically short acute phase associated with febrile illness and rashes, excruciating pain in multiple joints represents the most severe consequence of a CHIKV or MAYV infection in humans. The arthritis-like pain often manifests itself during the acute phase of the infection, but can persist in a subgroup of patients for months to years [4–6]. Symptoms cause a severe loss of quality of life and high economic costs, which is a burden especially for low-income countries [7]. The underlying pathophysiology of the chronic symptoms remains largely unclear, but appears to associate with circulating IL-6 [8] and IL-12 [9]. Furthermore, it may involve persisting viral RNA [9, 10], although this scenario has been debated [11].

Multiple studies on alphaviruses in immortalized model cell lines and *in vivo* in immunodeficient mice have provided valuable information on key aspects of CHIKV and MAYV tropism and replication, including host factors for entry and replication [12, 13], the impact of mutations in the viral glycoproteins on cell entry [14], and cellular restriction factors acting against CHIKV and other alphaviruses [15, 16]. Additionally, studies investigating immune responses to infection have demonstrated that CHIKV nsP2 counteracts host immunity by blocking nuclear translocation of STAT1 [17, 18] and inducing a host transcriptional shutdown [19, 20]. However, the relevance of these and potentially additional immunity-subverting mechanisms in infected patients remains unclear. *In vivo* studies in mice, though recapitulating both innate and adaptive immune responses, require a type I interferon (IFN)-deficient background, neglecting the impact of type I IFN-mediated antiviral responses [21]. However, type I IFN induced in and acting on nonhematopoietic cells appears to be essential for the control and early clearance of CHIKV *in vivo* [22–24]. Therefore, these systems do not fully recapitulate the cellular environment of human primary cells and tissues that are targeted by CHIKV and MAYV *in vivo*. Primary human cells have been used sporadically, but only few studies properly characterized their unique properties [25–27]. Here, we perform an in depth-characterization of primary human synovial fibroblasts as an *ex vivo* model of CHIKV and MAYV infection. Synovial fibroblasts have been described to be a key driver for rheumatoid arthritis by facilitating proinflammatory processes and stimulating the degradation of cartilage [28, 29]. Here, we establish synovial fibroblasts as being fully susceptible and permissive to CHIKV and MAYV infection. Using bulk and single-cell approaches, we identified cell-intrinsic immune responses that were most pronounced in abortively infected bystander cells, suggestive of effective viral antagonism of innate immunity in productively infected cells.

## Material and Methods

### Cells and Viruses

Human osteosarcoma U2OS cells (a kind gift from T. Stradal, Hanover), human HEK293T cells (a kind gift from J. Bohne, Hanover), human foreskin fibroblast HFF-1 cells (ATCC SCRC-1041), human HL116 cells (a kind gift from Sandra Pellegrini, Institut Pasteur, France [30]), and hamster BHK-21 cells (ATCC CCL-10) were grown in Dulbecco’s modified Eagle’s medium - high glucose (DMEM, Sigma-Aldrich D5671) supplemented with 10 % heat-inactivated fetal bovine serum (FBS, Sigma-Aldrich F7524), 2mM L-Glutamine (Gibco 25030081), and 100 units/ml penicillin-streptomycin (Gibco 11548876). HL116 cell received 1X HAT supplement (Gibco 21060017) in addition. Primary human fibroblasts were obtained from synovial biopsies from donors suffering from osteoarthritis (osteroarthrosis synovial fibroblasts, OASF) or a non-arthritic background (healthy donor synovial fibroblasts, HSF), purified, and cultured as described before [31]. The local ethic committee (Justus-Liebig-University Giessen) approved the cooperative study (ethical vote IDs 66-08 and 74-05). All patients gave written informed consent. Mycoplasma testing was routinely performed and negative in all primary human cell cultures. After 2-4 passages of initial cultivation, cells were expanded and used for experiments in high glucose DMEM supplemented with 20 % FBS, 2 mM L-Glutamine, 100 units/ml penicillin-streptomycin, 1 % non-essential amino acids (Gibco 11140050), and 1 % sodium pyruvate (Gibco 11360070). The CHIKV LR2006-OPY 5’GFP and MAYV TRVL4675 5’GFP infectious clones expressing EGFP under the control of a subgenomic promotor (hereafter referred to as CHIKV and MAYV) have been described previously [32, 33]. Virus was produced by *in vitro*-transcription of and subsequent electroporation of RNA into BHK-21 cells. Virus-containing supernatant was collected, passaged once on BHK-21 cells and viral titers were determined by titration on HEK293T cells.

### Infection, Treatments, Transfections

EGFP expression as surrogate for productive CHIKV or MAYV infection was quantified on a BD FACSCalibur, FACSLyric or Accuri C6. For neutralization assays, virus-containing supernatants were pre-incubated for one hour with anti-CHIKV E2 antibody C9 (Integral Molecular C9, Lot INT MAB-003) at 1 μg/ml or with recombinant MXRA8-Fc (a kind gift from M. Diamond) at 150 ng/ml. Recombinant IFN-α2a (Roferon L03AB04, Roche) and IFN-λ1 (Peprotech 300-02L) was used where indicated. Transfections were performed using Lipofectamine2000 (Thermo Fisher 11668019) for plasmid DNA (pcDNA6 empty vector) or 5’triphosphate dsRNA (InvivoGen tlrl-3prna).

### Bulk RNA-Seq Analysis

RNA was extracted using the Promega Maxwell 16 with LEV simplyRNA Tissue Kits (Promega AS1270). RNA quality was assessed using the Agilent Bioanalyzer and appropriate samples were used for NGS library preparation with the NEBNext Ultra II Directional RNA kit (NEB E7760) and sequenced with 50 bp paired-end reads and 30 mio reads per sample on the Illumina HiSeq 2500. Data was analyzed with CLC Genomics Workbench 12 (QIAGEN) by mapping the human reads onto the hg19 reference genome scaffold (GCA_000001405.28). Unmapped reads not matching the human genome were subsequently mapped onto the CHIKV genome LR2006_OPY (DQ443544.2). For HSF, infection and analysis were performed similarly, but RNA was extracted with the Direct-Zol RNA MiniPrep Kit (Zymo Research R2051), NGS libraries were prepared with the TruSeq stranded mRNA kit (Illumina 20020594) and sequencing was performed on the Illumina NextSeq500 with 65 mio reads per sample. Biological process enrichment was analyzed by Gene Ontology [34, 35].

### Single-Cell RNA-Seq Analysis

Infected cells were trypsinized, debris was removed by filtration, and the suspension was adjusted to a final amount of ∼16,000 cells per lane to achieve the recovery of 10,000 cells per donor after partitioning into Gel-Beads in Emulsion (GEMs) according to the instructions for Chromium Next GEM Single Cell 3’ GEM, Library & Gel Bead Kit v3.1 provided by the manufacturer (10X Genomics PN-1000121). Polyadenylated mRNAs were tagged with unique 16 bp 10X barcodes and the 10 bp Unique Molecular Identifiers (UMIs), reverse transcribed and resulting cDNAs were bulk amplified. After enzymatic fragmentation and size selection, resulting double-stranded cDNA amplicons optimized for library construction were subjected to adaptor ligation and sample index PCRs needed for Illumina bridge amplification and sequencing. Single-cell libraries were quantified using Qubit (Thermo Fisher) and quality-controlled using the Bioanalyzer System (Agilent). Sequencing was performed on a HiSeq4000 device (Illumina) aiming for 175 mln reads per library (read1: 26 nucleotides, read2: 64 nucleotides). Data was analyzed using CellRanger v5.0 (10X Genomics) using human and CHIKV genome scaffolds as described above, and the R packages Seurat v4.0 [36] and DoRothEA v3.12 [37].

### Quantitative RT-PCR

RNA was extracted using the Promega Maxwell 16 with the LEV simplyRNA tissue kit (Promega AS1270), the Roche MagNAPure with the Cellular Total RNA Large Volume kit (Roche 05467535001), or the DirectZol RNA Mini kit (Zymo R2051). cDNA was prepared using dNTPs (Thermo Fisher R0181), random hexamers (Jena Bioscience PM-301) and M-MuLV reverse transcriptase (NEB M0253). For quantitative RT-PCR, specific Taqman probes and primers (Thermo Fisher 4331182) were used with TaqMan Universal PCR Master Mix (Applied Biosystems 4305719) or LightCycler© 480 Probes Master (Roche 04887301001). PCRs were performed on the Applied Biosystems ABI 7500 Fast or the Roche LightCycler 480 in technical triplicates.

### Flow Cytometry, Confocal and Live Cell Imaging

For flow cytometric analysis of protein expression, OASF were fixed in 4 % PFA (Carl Roth 4235.2), permeabilized in 0.1 % Triton-X (Invitrogen HFH10) and immunostained with antibodies against IFIT1 (Origene TA500948, clone OTI3G8), MX1/2 (Santa Cruz sc-47197), and IFITM3 (Abgent AP1153a) in combination with Alexa Fluor-647 conjugated antibodies against mouse-(Thermo Fisher A28181), rabbit- (Thermo Fisher A27040), or goat-IgG (Thermo Fisher A-21447). Flow was performed on a BD FACSCalibur or FACSLyric and analyzed with FlowJo v10. For immunofluorescence microscopy, OASF were seeded in 8-well μ-slides (ibidi 80826), fixed and permeabilized as described above, stained with antibodies against MXRA8 (biorbyt orb221523) with AlexaFluor647-conjugated secondary antibody (Thermo Fisher A28181), and counterstained with DAPI (Invitrogen D1306). For fluorescence microscopy and live cell imaging, cells were infected with CHIKV at an MOI of 10 and imaged with the Zeiss LSM800 Airyscan Confocal Microscope. Images were analyzed and merged using Zeiss ZEN Blue 3.0.

### Immunoblotting

Cell lysates were separated on 10 % acrylamide gels by SDS-PAGE and protein transferred to a 0.45 μm PVDF membrane (GE Healthcare 15259894) using the BioRad TransBlot Turbo system. Expression was detected using primary antibodies detecting MXRA8 (biorbyt orb221523), FHL1 (R&D Systems MAB5938), IFITM3 (Abgent AP1153a), MX2 (Santa Cruz sc-47197), ISG15 (Santa Cruz sc-166755), and α-Tubulin (Cell Signaling Technology 2144S) and appropriate secondary IRDye antibodies. CHIKV proteins were detected using anti-CHIKV antiserum (IBT Bioservices Cat #01-0008 Lot #1703002). Fluorescence was detected and quantified using the LI-COR Odyssey Fc system.

### Measurement of IL-6 and Bioactive IFN

Anti-IL-6 ELISA (BioLegend 430504) was performed according to manufacturer’s protocols. Briefly, plates were coated with capture antibodies and incubated with diluted supernatant from CHIKV- or mock-infected cell cultures. Detection antibody and substrate were added and the OD measured with the Tecan Sunrise microplate reader. Concentrations were then calculated the concentration according to a standard curve measured on the same plate. Bioactive type I IFN was quantified by incubating supernatant from CHIKV-infected cells on HL116 cells harboring a firefly luciferase gene under the control of an IFN-sensitive promotor. After six h, cells were lysed, incubated with luciferase substrate solution (Promega E1500), and luciferase activity was quantified with the BioTek Synergy HTX microplate reader.

### Data and Code Availability

RNA-seq and single-cell RNA-seq datasets are available at the NCBI GEO database under the accession number GSE152782 and GSE176361, respectively. All generated code is available at https://github.com/GoffinetLab/CHIKV_scRNAseq-fibroblast.

### Data Presentation and Statistical Analysis

If not stated otherwise, bars and symbols show the arithmetic mean of indicated amount of repetitions. Error bars indicate S.D. from at least three or S.E.M. from the indicated amount of individual experiments. Statistical analysis was performed using CLC Workbench for RNA-seq and GraphPad Prism 8.3.0 for all other analysis. Unpaired t-tests were applied with assumed equal standard deviation when comparing results obtained in the same cell line and Mann-Whitney-U-tests when comparing between cell lines or between cell lines and primary cells. For treatment analysis, ratio paired t-tests were applied. For IC50 calculation, nonlinear fit curves with variable slopes were calculated. FDR correction was applied for RNA-seq analysis and Bonferroni correction for Gene Ontology analysis. *P* values <0.05 were considered significant (*), <0.01 very significant (**), <0.001 highly significant (***); < 0.0001 extremely significant, n.s. = not significant (≥0.05).

## Results

### Osteoarthritic fibroblasts are susceptible and permissive to CHIKV and MAYV infection

First, we examined the ability of primary human synovial fibroblasts to support the entire CHIKV and MAYV replication cycle. Therefore, we infected synovial fibroblasts obtained from osteoarthritic patients (OASF) and from patients with a non-arthritic background (HSF) with CHIKV strain LR2006-OPY or MAYV strain TRVL7546 expressing EGFP under the control of a second subgenomic promotor. 24 hours post-infection, the proportion of EGFP-positive cells ranged between 4 and 24.5 % for CHIKV and between 8.5 and 39 % for MAYV and did not differ between fibroblast types (Fig. 1A). At the same time point, supernatants of both OASF and HSF displayed CHIKV titers of 1.6-8.8x10^5^ infectious particles per ml and MAVV titers of 0.12-2.75x10^5^ infection particles per ml, with significantly higher titers produced by OASF. At 48 hours post-infection, CHIKV titers produced by HSF did not further increase, whereas the titers produced by OASF reached up to 1.5x10^7^ infectious particles per ml (Fig. 1B, left panel), suggesting slightly higher virus production and/or viral spread in OASF as compared to HSF. MAYV titers did not significantly increase in OASF or HSF at 48 hours post-infection (Fig. 1B, right panel).

**Figure 1.**
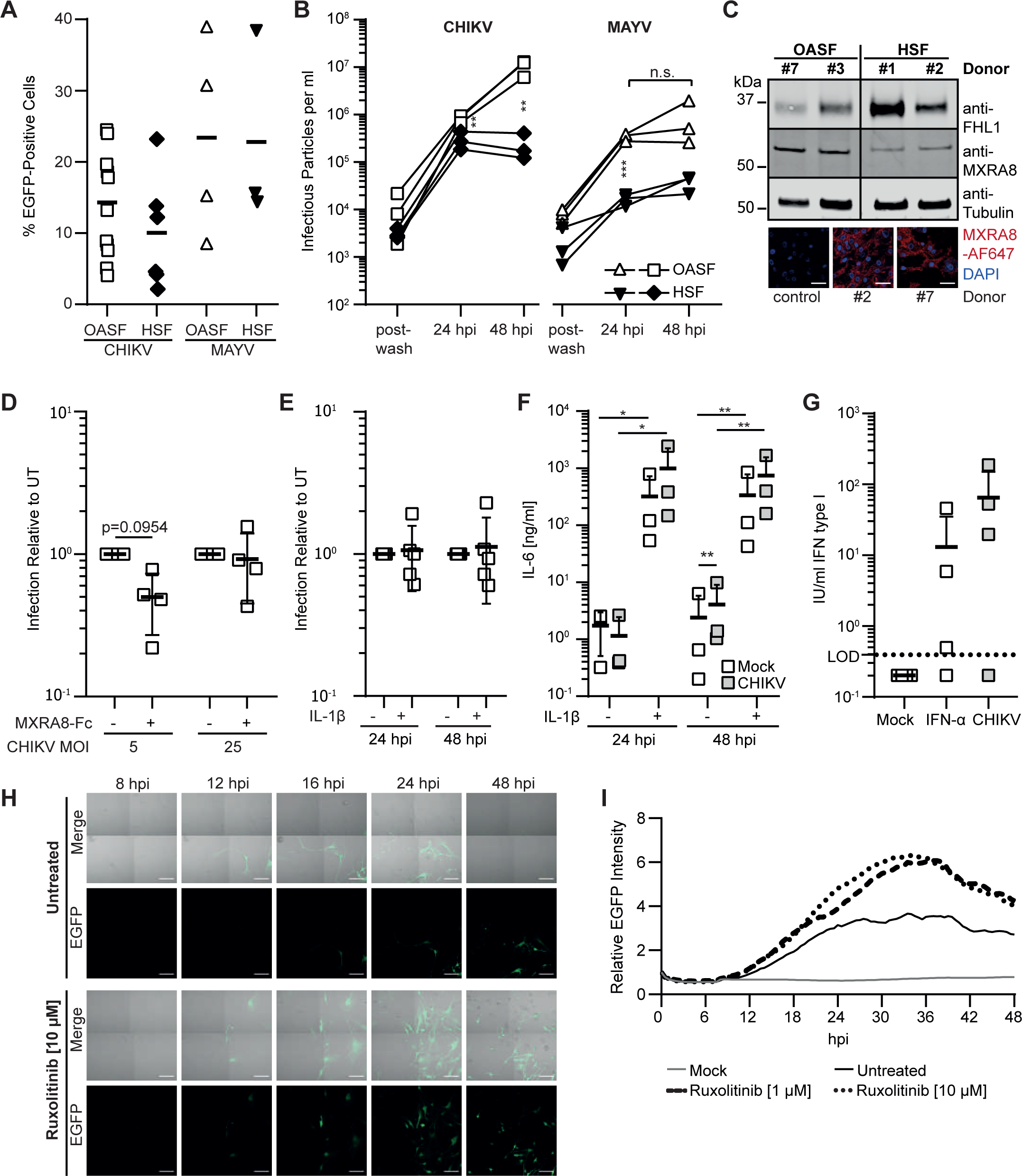
Osteoarthritis fibroblasts are susceptible and permissive to CHIKV infection. (**A**) OASF or HSF were infected with 5’EGFP-CHIKV or -MAYV (MOI 10). 24 hours post-infection, the percentage of EGFP-positive cells was quantified by flow cytometry (n = 3-12). (**B**) Supernatants of CHIKV- and MAYV-infected OASF or HSF were collected at 24 and 48 hours post-infection, and titers were determined by analyzing EGFP expression at 24 hours post-infection of HEK293T cells. For background controls (post-wash), samples were taken after one hour of virus inoculation and subsequent washing (n = 3). (**C**) Uninfected OASF and HSF were analyzed for MXRA8 and FHL1 expression by immunoblotting (n = 4-6) and for MXRA8 expression by immunofluorescence. Scale bar = 50 μm (n = 3, representative images shown). (**D**) OASF were infected with 5’EGFP-CHIKV at the indicated MOIs upon treatment of the virus with MXRA8-Fc recombinant protein or mock treatment. At 24 hours post-infection, cells were analyzed for EGFP expression (n = 4). (**E**) OASF were stimulated with IL-1β at 10 ng/ml for 16 hours and subsequently infected with CHIKV (MOI 10) in the presence of IL-1β. Mock-stimulated OASF were infected as a control. At 24 and 48 hours post-infection, cells were analyzed for EGFP expression and (**F**) supernatant was collected and analyzed for IL-6 secretion by ELISA (n = 3). (**G**) OASF were infected with 5’EGFP-CHIKV or treated with 200 IU/ml IFN-α. 24 h later, supernatants of the infected cells were incubated on HL116 reporter cells to quantify secreted bioactive type I IFN (n = 4). (**H**) OASF were infected with 5’EGFP-CHIKV (MOI 10) in the presence or absence of 1 or 10 μM Ruxolitinib or mock-infected. Infection was recorded by live-cell imaging and representative images for untreated and 10 μM Ruxolitinib-treated cells are shown. Scale bar = 100 μm. (**I**) Infected cells recorded by live-cell imaging in G were analyzed for EGFP intensity using ImageJ (n = 3).

Susceptibility of cells to CHIKV infection is enhanced by the attachment factor MXRA8 [12] and the cytosolic protein FHL-1 is essential for CHIKV genome replication [13]. We confirmed expression of these two cellular cofactors in OASF and HSF by immunoblotting and/or immunofluorescence (Fig. 1C). We assessed the functional relevance of the MXRA8 attachment factor using a soluble MXRA8-Fc fusion protein, which blocks the binding site on the E1-E2 glycoprotein complex on the virus surface [12, 38]. At a low MOI, MXRA8-Fc-preincubated CHIKV was 50 % less infectious to synovial fibroblasts, and this inhibition was abolished when saturating amounts of infectious virus particles were used (Fig. 1D), indicating that endogenous MXRA8 contributes, at least partially, to CHIKV entry in OASF.

Subsequently, we investigated whether IL-1β-mediated activation of synovial fibroblasts, a hallmark of rheumatoid arthritis [39–41], modulates their susceptibility to CHIKV infection. Treatment with IL-1β, did not alter the percentage of EGFP-positive cells upon CHIKV challenge (Fig. 1E), while readily inducing IL-6 secretion (Fig. 1F). Conversely, CHIKV infection only very mildly, if at all, enhanced IL-1β-induced IL-6 secretion (Fig. 1F). These data suggest that CHIKV infection of synovial fibroblasts neither induces nor modulates IL-6 secretion, arguing against their activation.

To determine the importance of IFN-mediated antiviral immunity in this primary cell system, we analyzed the secretion of type I IFN upon CHIKV infection, which reached higher levels than after prestimulation with IFN-α (Fig. 1G). Additionally, we monitored the CHIKV infection in the absence or presence of the JAK/STAT inhibitor Ruxolitinib. Using live-cell imaging, we documented the increase in EGFP-positive cells between ten and 48 hours post-infection, which progressed faster in Ruxolitinib-treated cultures, with an onset of cytopathic effects observed after 24 hours in all infected cultures (Fig. 1H, Suppl. Mov. 1). Analysis of the EGFP intensity in each frame over time confirmed the higher expression of EGFP in Ruxolitinib-treated cultures (Fig. 1I, Suppl. Mov. 2). Overall, these experiments establish the susceptibility and permissiveness of synovial fibroblasts to CHIKV and MAYV infection and their expression of important cellular cofactors. Furthermore, we show an absence of interconnection between IL1-β activation and susceptibility to CHIKV infection, and restriction of infection through JAK/STAT-mediated innate immunity.

### CHIKV infection provokes a strong cell-intrinsic immune response in OASF

Next, we performed RNA-seq analysis on OASF and HSF that had been infected with CHIKV in the presence or absence of the glycoprotein E2-binding, neutralizing antibody C9 [42], and on mock-infected cells. C9 pre-treatment resulted in potent inhibition of the infection by on average 16-fold (Fig. 2A). Upon infection, expression of numerous IFN-stimulated genes (ISGs) was induced at the protein level in a C9 treatment-sensitive manner, including IFITM3, ISG15, and MX2. As expected, production of the viral E1-E2 and capsid proteins was detectable specifically in CHIKV-infected, but not in cells exposed to C9-pretreated virus (Fig. 2B). Global transcriptional profiling by RNA-seq revealed 992 (OASF) and 1221 (HSF) upregulated genes as well as 99 (OASF) and 353 (HSF) downregulated genes in CHIKV-infected cells 24 hours post-infection as compared to uninfected cells (Fig. 2C). Uninfected cells and cells exposed to C9-treated virus shared a similar profile (data not shown). A high similarity of the gene expression profile of uninfected OASF and HSF (R^2^=0.9086) argues against a potential transcriptional predisposition that could have exerted a rheumatoid arthritis-related gene expression profile or a broad proinflammatory activation (Fig. S1A). Uninfected OASF and HSF differed in genes involved in organ development and cellular regulatory processes, and not inflammatory or antiviral processes (Fig. S1B). Additionally, the transcriptional profile in infected OASF and HSF was very similar (R^2^=0.9085, Fig. S1C-D), with an equivalently strong upregulation of a set of prototypic inflammation and arthritis-related genes which we defined for further analysis (R^2^=0.8202, Fig. 2D). Interestingly, the number of genes significantly up- and downregulated upon infection was 1.23-fold and 3.57-fold higher in HSF compared to OASF, respectively, but 55.4% of upregulated genes from both groups overlapped (Fig. 2E). Most of the prototypic antiviral and proinflammatory genes were highly upregulated in infected cells, demonstrating a broad and strong activation of antiviral immune responses in cells from four different donors with no statistically significant deviation in the magnitude of induction (Fig. 2F, left panel). Upregulation of *IFNB* and *IFNL1*, *IFNL2*, and *IFNL3* expression was statistically significant but low in magnitude, with almost no *IFNA* mRNA detectable. Expression of arthritis-associated genes, including genes encoding immune cell chemoattractants (*CXCL5, IL8, CD13, RANTES/CCL5*), matrix-metalloproteases (*MMP3, -9, -14, ADAMTS5*) and genes commonly expressed by fibroblasts in rheumatoid arthritis (*FGF2, PDPN, NGF, FAP*), was not grossly altered in CHIKV-infected cells. Exceptions were a strong CHIKV-induced upregulation of *RANTES/CCL5* in both OASF and HSF and *IL8* in HSF (Fig. 2F, right panel). mRNAs for all IFN receptors were detectable and stable with exception of *IFNLR1*, whose expression was upregulated upon CHIKV infection (Fig. S1E). Established host factors for CHIKV as well as fibroblast marker genes and cellular housekeeping genes were not quantitatively altered in their expression. Virtual absence of expression of monocyte/macrophage lineage-specific genes excluded the possibility of a contamination of the fibroblast culture with macrophages, which occasionally has been reported in early passages of *ex vivo-*cultured synovial fibroblasts [31] (Fig. S1E). Conclusively, OASF and HSF share similar basal and CHIKV infection-induced transcriptional profiles. Overall, CHIKV-infected OASF sense and react to productive CHIKV infection with the extensive upregulation of antiviral and proinflammatory ISGs. IFN expression itself was low at 24 hours post-infection, not excluding the possibility that it peaked transiently at earlier time points.

**Figure 2.**
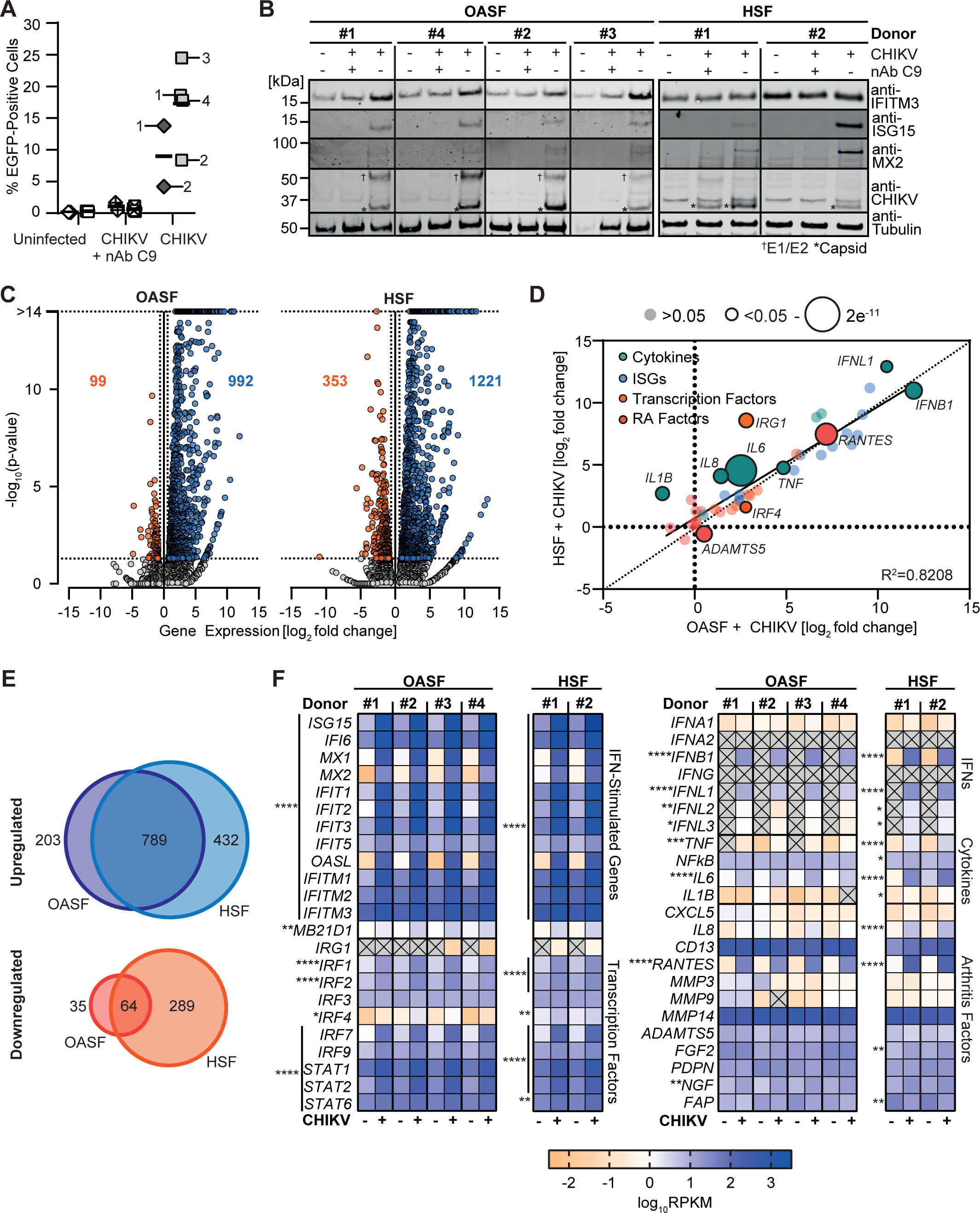
Productive CHIKV infection provokes a strong cell-intrinsic immune response in OASF and HSF. (**A**) OASF were infected with 5’EGFP-CHIKV at an MOI of 10 in the presence or absence of the anti-E2 antibody C9 and the percentage of EGFP-positive cells was measured by flow cytometry (OASF: squares, HSF: diamonds. The infected samples are marked with their respective donor number). (**B**) Selected proteins of cells infected in A were analyzed by immunoblotting. (**C-F**) RNA from cells infected in A was extracted and subjected to RNA-seq (n = 4). (**C**) Analysis of up- and downregulated genes in CHIKV-infected samples compared to mock. Dotted lines indicate cutoff for <1.5 fold regulation and a p-value of >0.05. (**D**) Visualization of the fold change induction of indicated genes in CHIKV-infected OASF and HSF. Average fold change (log2) values for infected OASF are plotted on the x-axis, with corresponding values from infected HSF plotted on the y-axis. R^2^ value and regression line for the comparison are inset, dot sizes indicate significance. (**E**) Overlap of significantly (FDR-p <0.05) up- and downregulated genes in infected OASF and HSF. Numbers of genes up- or downregulated in either OASF or HSF only, or in both cell-types, are indicated. (**F**) Heatmaps of selected gene expression profiles related to innate immune responses (left) or to secreted proinflammatory mediators and arthritis-connected genes (right) in uninfected or CHIKV-infected cells.

### High viral RNA levels in cells of infected cultures with an excess replication of the viral structural subgenome

We noticed very little inter-donor variation regarding the distribution of identified viral reads along the viral genome. The 5’ region of the genome, encoding the non-structural CHIKV proteins, was replicated to a lower extent than the 3’, 26S subgenomic promoter-driven, structural protein-encoding genomic region. Interestingly, this differential abundance of 5’and 3’ reads was also detected in cultures inoculated with C9-neutralized virus, suggesting infection in a small number of cells (Fig. 3A). Overall, the 26S subgenomic viral RNA was 5.3-fold more abundant than nonstructural subgenomes (Fig. 3B). 18-54 % and 17-44 % of the total reads in productively infected OASF and HSF, respectively, were attributed to the CHIKV genome (Fig. 3C, D). In summary, our analysis revealed efficient replication of the CHIKV genome in infected fibroblasts with an excess of structural protein-encoding subgenomic RNA.

**Figure 3.**
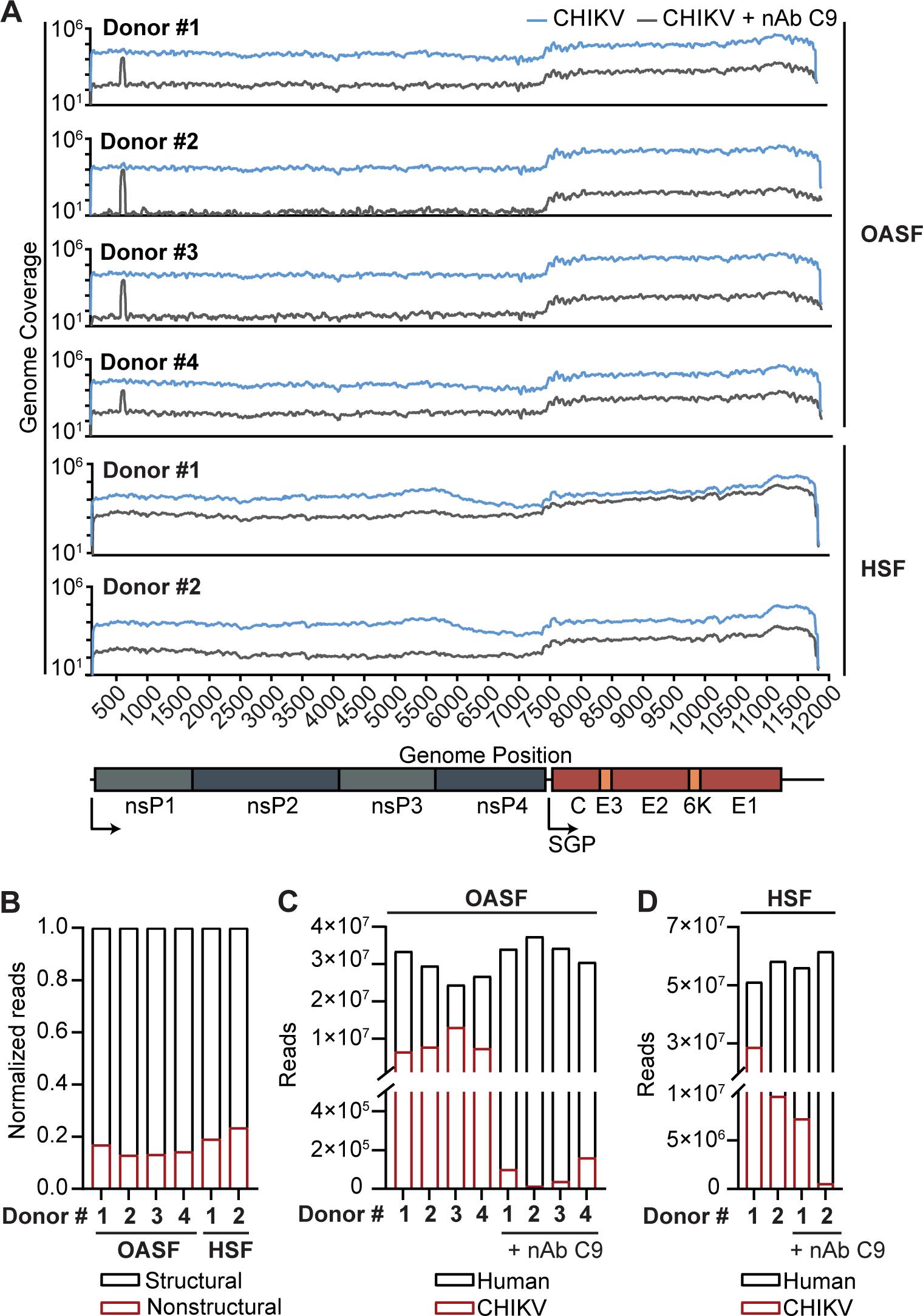
The CHIKV genome is replicated to a high degree with a strong bias towards the structural subgenome in infected OASF. (**A**) NGS reads attributed to each individual position in the CHIKV genome plotted for cells infected with CHIKV in the presence or absence of neutralizing antibody (nAb). SGP: subgenomic promotor. (**B**) Normalized amount of reads attributed to the structural and nonstructural part of the CHIKV genome in CHIKV-infected OASF and HSF. (**C**) Number of NGS reads attributed to the human or CHIKV reference genome in CHIKV or neutralizing antibody-treated CHIKV infected OASF or (**D**) HSF.

### Exogenous IFN administration provokes higher immune responses and leads to improved protection from infection in primary fibroblasts than in commonly used cell lines

CHIKV and MAYV infection rates in OASF did not increase after 24 hours post-infection (Fig. 4A), and we suspected this to be the result of the strong immune activation and subsequent IFN signaling. The commonly used osteosarcoma cell line U2OS was more susceptible, while the immortalized fibroblast cells line HFF-1 displayed reduced susceptibility to alphaviral infection (Fig. 4A). OASF exhibited strong induction of *IFIT1* and *MX2* CHIKV infection, which exceeded those mounted by U2OS and HFF-1 cells at both 24 and 48 hours post-infection by 15- to 150-fold. MAYV infection-provoked ISG responses in OASF were inferior to those induced by CHIKV, despite similar percentages of infected cells (Fig. 4B). Contrasting the cell system-specific magnitude of gene expression upon CHIKV infection, both OASF and cell lines shared similar responsiveness to 5’-triphosphate dsRNA (5-ppp-RNA) transfection, which exclusively stimulates the RNA sensor RIG-I [43], the main sensor of CHIKV RNA in infected cells [44], and plasmid DNA transfection (Fig. S2A).

**Figure 4.**
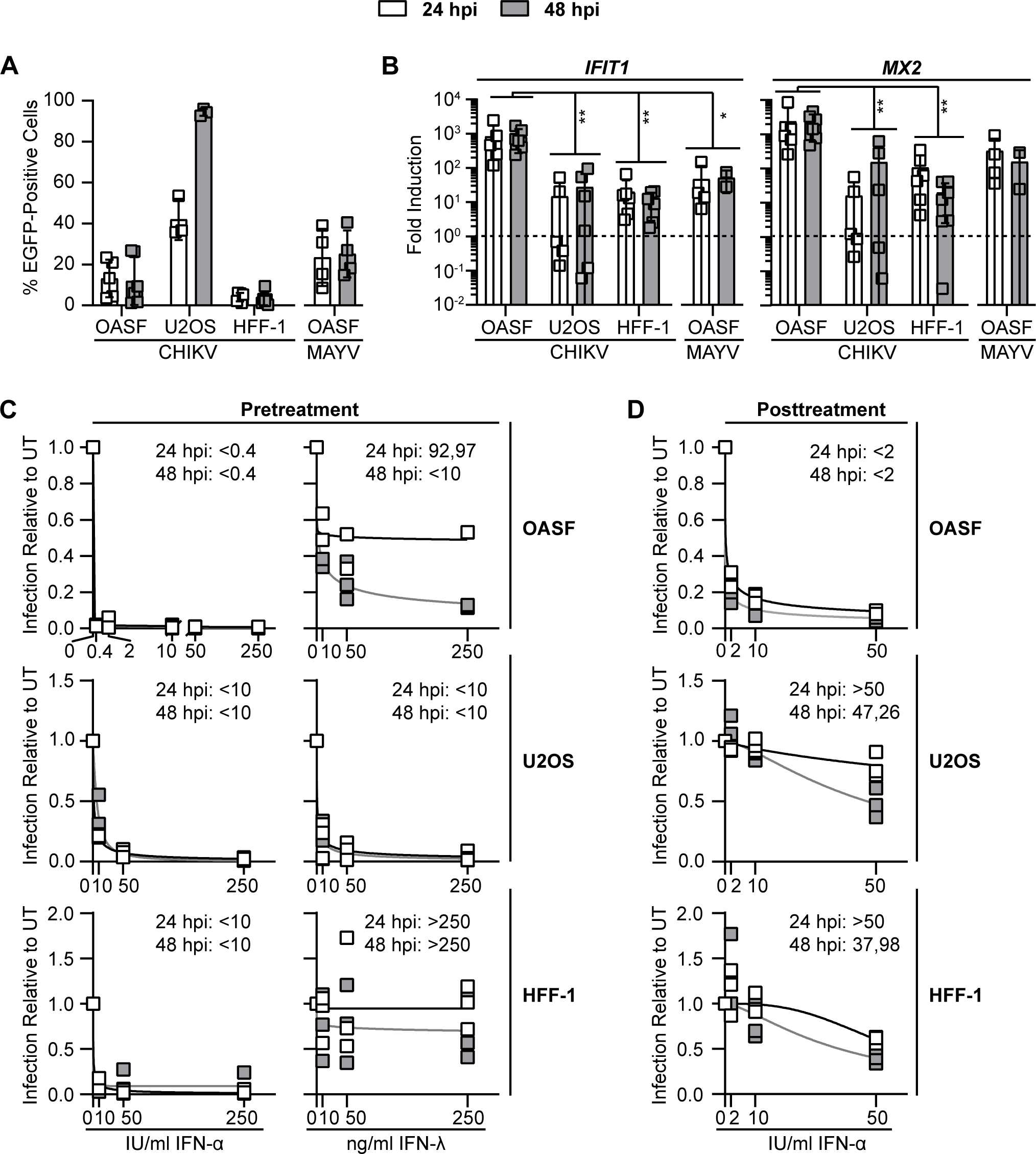
OASF react to CHIKV infection more strongly than commonly used cell lines and can potently inhibit viral infection after IFN treatment. (**A**) OASF and HFF-1 cells were infected with 5’EGFP-CHIKV at an MOI of 10, U2OS cells were infected at an MOI of 0.5. EGFP-positive cells were quantified at 24 and 48 hours post-infection by flow cytometry (n = 3-6). (**B**) Cells infected in (**A**) were analyzed for expression of *IFIT1* and *MX2* mRNA at 24 and 48 hours post-infection by quantitative RT-PCR (n = 3-6). (**C**) Cells were treated with IFN-α or -λ for 48 h before infection with 5’EGFP-CHIKV (OASF and HFF-1: MOI 10; U2OS: MOI 0.5) in the continuous presence of IFN. Inset numbers indicate IC50 values for each timepoint. (**D**) Cells were infected with 5’-EGFP CHIKV (OASF and HFF-1: MOI 10; U2OS: MOI 0.5) and IFN-α was added four hours post-infection. 24 and 48 hours post-infection, EGFP-positive cells were quantified by flow cytometry. Inset numbers indicate IC50 values for each timepoint. UT: untreated, IU: international units (n = 3 for all experiments)

Next, we tested the cellś ability to respond to exogenous type I and III IFNs, which play a crucial role in limiting virus infection and protecting the host [45–47]. We stimulated OASF individually with a range of IFN-α2 and -λ concentrations at 48 hours prior to infection. At all investigated concentrations, even the lowest dose, of IFN-α induced a potent upregulation of *IFIT1* and *MX2* (Fig. S2B), and almost completely inhibited CHIKV infection (Fig. 4C). In contrast, IFN-λ induced lower ISG expression levels (Fig. S2B), and inhibited infection less efficiently (Fig. 4C). Although less effective than in OASF, IFN-α restricted CHIKV infection both in U2OS and HFF-1 cells, while IFN-λ pre-treatment was more potent in U2OS cells than in OASF, and ineffective in HFF-1 cells (Fig. 4C). These antiviral activities were largely consistent with the respective degree of ISG expression at the time point of infection (Fig. S2B). IFN-α and -λ induced expression of *IFIT1* and *MX2* was higher in U2OS cells than in HFF-1. We next investigated the sensitivity of CHIKV infection to IFN when applied four hours post-infection. In this set-up, IFN-α still displayed a clear, though less potent antiviral activity when compared to the pre-treatment setting (Fig. 4D). In contrast, treatment of both immortalized cell lines with IFN-α post-infection was very ineffective (Fig. 4D).

Interestingly, in all three cells systems, a preceding CHIKV infection did not antagonize IFN-mediated induction of ISGs, and led to expression levels of *IFIT1* and *MX2* exceeding those induced by IFN-α alone (Fig. S2C). Overall, the data suggest a stronger sensitivity of OASF to IFN-α-induced immunity compared to commonly used immortalized cell lines. Most interestingly, and in striking contrast to the immortalized cell lines, OASF were unique in their ability to transform a post-infection treatment of IFN-α into a relatively potent antiviral program. Collectively, these data uncover crucial differences between primary synovial fibroblasts and widely used immortalized cell lines regarding their cell-intrinsic innate response to infection and their sensitivity to exogenous IFNs.

### Virus-inclusive single-cell sequencing reveals a switch from induction to repression of immune responses depending on a threshold level of viral RNA in infected cells

Finally, we asked how the cell-intrinsic defenses correlate with the amounts of viral RNA within cells of a given infected culture by analyzing infected OASF for their expression of antiviral proteins using flow cytometry. As expected, expression of IFIT1, IFITM3 and MX1/2 was enhanced in OASF upon IFN-α treatment (Fig. 5A). Interestingly, these proteins were expressed at even higher levels in EGFP-negative cells of CHIKV-infected cultures, while the productively infected, EGFP-positive cells displayed markedly reduced expression levels of these factors (Fig. 5A).

**Figure 5.**
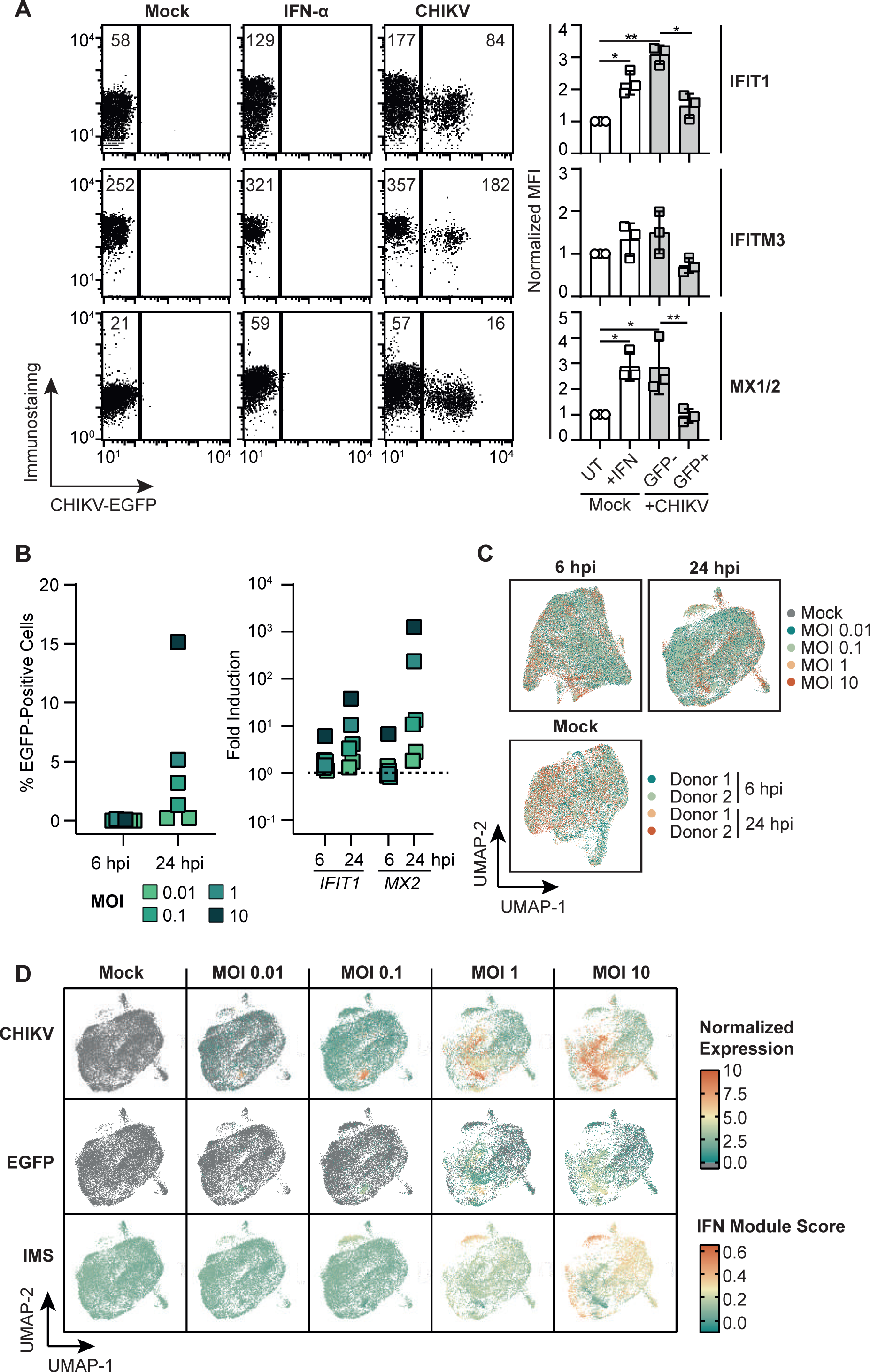
Reduced induction of antiviral protein and gene expression in productively infected cells. (**A**) OASF were infected with 5’EGFP-CHIKV (MOI 10) or treated with 100 IU/ml IFN-α and immunostained for IFIT1, IFITM3, and MX1/2 24 hours post-infection. Numbers in the dot plots indicate mean fluorescence intensities (MFI) of one representative experiment, and the bar diagram shows quantification of three individual experiments. (**B**) OASF were infected with 5’EGFP-CHIKV at indicated MOIs. Six and 24 hours post-infection, EGFP-positive cells were quantified by flow cytometry (left panel), and cells were analyzed for expression of *IFIT1* and *MX2* mRNA (right panel, n = 6). (**C**) Using OASF infected with 5’EGFP-CHIKV, single-cell RNA-sequencing was conducted and UMAP visualizations for sample overlapping after integration are shown. (**D**) UMAP projections from infected OASF (24 hours post-infection) indicate the abundance of CHIKV 3’ end reads, EGFP 3’ end reads, and IFN signaling gene expression as calculated by IMS.

Since absence of EGFP expression does not necessarily exclude the presence of viral, potentially abortive RNA, we performed virus-inclusive single-cell RNA-seq to establish potential correlations of the quantity of viral RNA and a specific cellular transcriptional profile. To this end, we analyzed infected OASF infected at escalating MOIs. No EGFP-positive cells were detectable at six hours post-infection by flow cytometry (Fig. 5B, left panel). In contrast, 24 hours post-infection, the reporter was expressed in an MOI-dependent fashion, ranging from virtually 0% to 15% (Fig. 5B, left panel). *IFIT1* and *MX2* mRNA expression was largely proportional to EGFP expression (Fig, 5B, right panel).

Single-cell (sc) RNA-seq of the very same cells showed very little inter-donor variability, and we merged data from both donors throughout the rest of the analysis (Fig. 5C). In order to identify potential correlations of viral RNA abundance and the cellular transcription profile, we compared the expression of CHIKV RNA to expression of 203 IFN signaling genes listed in the REACTOME database (identifier R-HSA-913531, Table 1). For each cell, the expression of this collection of genes was summarized using Seurat’s AddModuleScore function. Briefly, this summarizes the expression of a select group of genes by normalizing the aggregate expression to a randomly selected, non-overlapping subset of genes and scores each cell based on its expression of genes in this module, creating a module score (IFN Module Score, IMS). 24 hours post-infection, most identified CHIKV reads corresponded to the 3’ end of the genome, along with a minor number of reads mapping to the 3’ end of *EGFP*, which is expressed as a subgenomic RNA in infected cells (Fig. S3A). As expected for mock-infected cells, CHIKV reads were undetectable, and IFN signaling genes were expressed at basal levels, as calculated by the IMS. CHIKV RNA abundance per cell increased in an MOI-dependent manner, however susceptibility to infection was unequally distributed over individual cells, and a subset of cells displayed a higher susceptibility than others, as reflected by a high percentage of reads attributed to the viral genome (Fig. S3B). Most interestingly, IFN signaling genes appeared to be induced predominantly in cells displaying low CHIKV gene expression. *Vice versa*, CHIKV RNA-positive cells maintained basal or reduced expression of IFN signaling related genes (Fig. 5D). Of note, six hours post-infection, CHIKV expression was low and antiviral responses as presented by the IMS was were largely absent at low MOIs, while individual ISGs were induced at higher MOIs (Fig. S3C, D). As opposed to the induction of IFN signaling genes, known CHIKV cofactors *MXRA8*, *FHL1*, and the fibroblast marker genes *VIM* and *COL3A1* were broadly and stably expressed under all experimental conditions. Surprisingly, *FURIN*, encoding the cellular protease considered important for viral polyprotein cleavage, was detectable only in a minority of cells (Fig. S4).

**Table 1:**

**Pott et al., Table 1**
|  |  |  |  |  |
| --- | --- | --- | --- | --- |
| AAAS | HLA-DRB3 | ISG15 | PLCG1 | TRIM68 |
| ABCE1 | HLA-DRB4 | ISG20 | PML | TRIM8 |
| ADAR | HLA-DRB5 | JAK1 | POM121 | TYK2 |
| ARIH1 | HLA-E | JAK2 | POM121C | UBA52 |
| B2M | HLA-F | KPNA1 | PPM1B | UBA7 |
| BST2 | HLA-G | KPNA2 | PRKCD | UBB |
| CAMK2A | HLA-H | KPNA3 | PSMB8 | UBC |
| CAMK2B | ICAM1 | KPNA4 | PTAFR | UBE2E1 |
| CAMK2D | IFI27 | KPNA5 | PTPN1 | UBE2L6 |
| CAMK2G | IFI30 | KPNA7 | PTPN11 | UBE2N |
| CD44 | IFI35 | KPNB1 | PTPN2 | USP18 |
| CIITA | IFI6 | MAPK3 | PTPN6 | USP41 |
| DDX58 | IFIT1 | MID1 | RAE1 | VCAM1 |
| EGR1 | IFIT2 | MT2A | RANBP2 | XAF1 |
| EIF2AK2 | IFIT3 | MX1 | RNASEL |  |
| EIF4A1 | IFITM1 | MX2 | RPS27A |  |
| EIF4A2 | IFITM2 | NCAM1 | RSAD2 |  |
| EIF4A3 | IFITM3 | NDC1 | SAMHD1 |  |
| EIF4E | IFNA1 | NEDD4 | SEC13 |  |
| EIF4E2 | IFNA10 | NUP107 | SEH1L |  |
| EIF4E3 | IFNA13 | NUP133 | SOCS1 |  |
| EIF4G1 | IFNA14 | NUP153 | SOCS3 |  |
| EIF4G2 | IFNA16 | NUP155 | SP100 |  |
| EIF4G3 | IFNA17 | NUP160 | STAT1 |  |
| FCGR1A | IFNA2 | NUP188 | STAT2 |  |
| FCGR1B | IFNA21 | NUP205 | SUMO1 |  |
| FLNA | IFNA4 | NUP210 | TPR |  |
| FLNB | IFNA5 | NUP214 | TRIM10 |  |
| GBP1 | IFNA6 | NUP35 | TRIM14 |  |
| GBP2 | IFNA7 | NUP37 | TRIM17 |  |
| GBP3 | IFNA8 | NUP42 | TRIM2 |  |
| GBP4 | IFNAR1 | NUP43 | TRIM21 |  |
| GBP5 | IFNAR2 | NUP50 | TRIM22 |  |
| GBP6 | IFNB1 | NUP54 | TRIM25 |  |
| GBP7 | IFNG | NUP58 | TRIM26 |  |
| HERC5 | IFNGR1 | NUP62 | TRIM29 |  |
| HLA-A | IFNGR2 | NUP85 | TRIM3 |  |
| HLA-B | IP6K2 | NUP88 | TRIM31 |  |
| HLA-C | IRF1 | NUP93 | TRIM34 |  |
| HLA-DPA1 | IRF2 | NUP98 | TRIM35 |  |
| HLA-DPB1 | IRF3 | OAS1 | TRIM38 |  |
| HLA-DQA1 | IRF4 | OAS2 | TRIM45 |  |
| HLA-DQA2 | IRF5 | OAS3 | TRIM46 |  |
| HLA-DQB1 | IRF6 | OASL | TRIM48 |  |
| HLA-DQB2 | IRF7 | PDE12 | TRIM5 |  |
| HLA-DRA | IRF8 | PIAS1 | TRIM6 |  |
| HLA-DRB1 | IRF9 | PIN1 | TRIM62 |  |

### Correlating viral and cellular gene expression reveals a selective suppression of transcription factor and ISG expression

In order to quantify expression of IFN signaling genes according to viral RNA abundance, we divided cells into three groups: cells without detectable viral RNA expression (bystander), cells displaying low amounts of viral RNA (low) and cells displaying high levels of viral RNA (high) (Fig. 6A). Mirroring our initial observations (Fig. 5), we detected a significantly lower IMS in high cells when compared to low or bystander cells of the identical culture (Fig. S5A). Six hours post-infection, differential expression of non-ISGs was very modest between bystander and viral RNA-positive cells, while it was clearly more pronounced 24 hours post-infection (Fig. S5B). In contrast, over 250 ISGs, including *ISG15*, *IFIT1*, *MX2*, *IFITM3*, *MX1*, and *IFI6*, were upregulated in viral RNA-positive cells as compared to bystander cells at both investigated time points. Individual comparisons of either low or high cells with bystander cells gave similar overall observations. However, at both investigated time points, no further upregulation of ISGs was detected in the high cells as compared to low cells, but rather a significant downregulation of three ISGs at 24 hours post-infection and six ISGs at six hours post-infection. This suggests either a loss of cellular transcription activity or a lowered stability of cellular RNA in cells containing high loads of CHIKV RNA (Fig. S5B).

**Figure 6.**
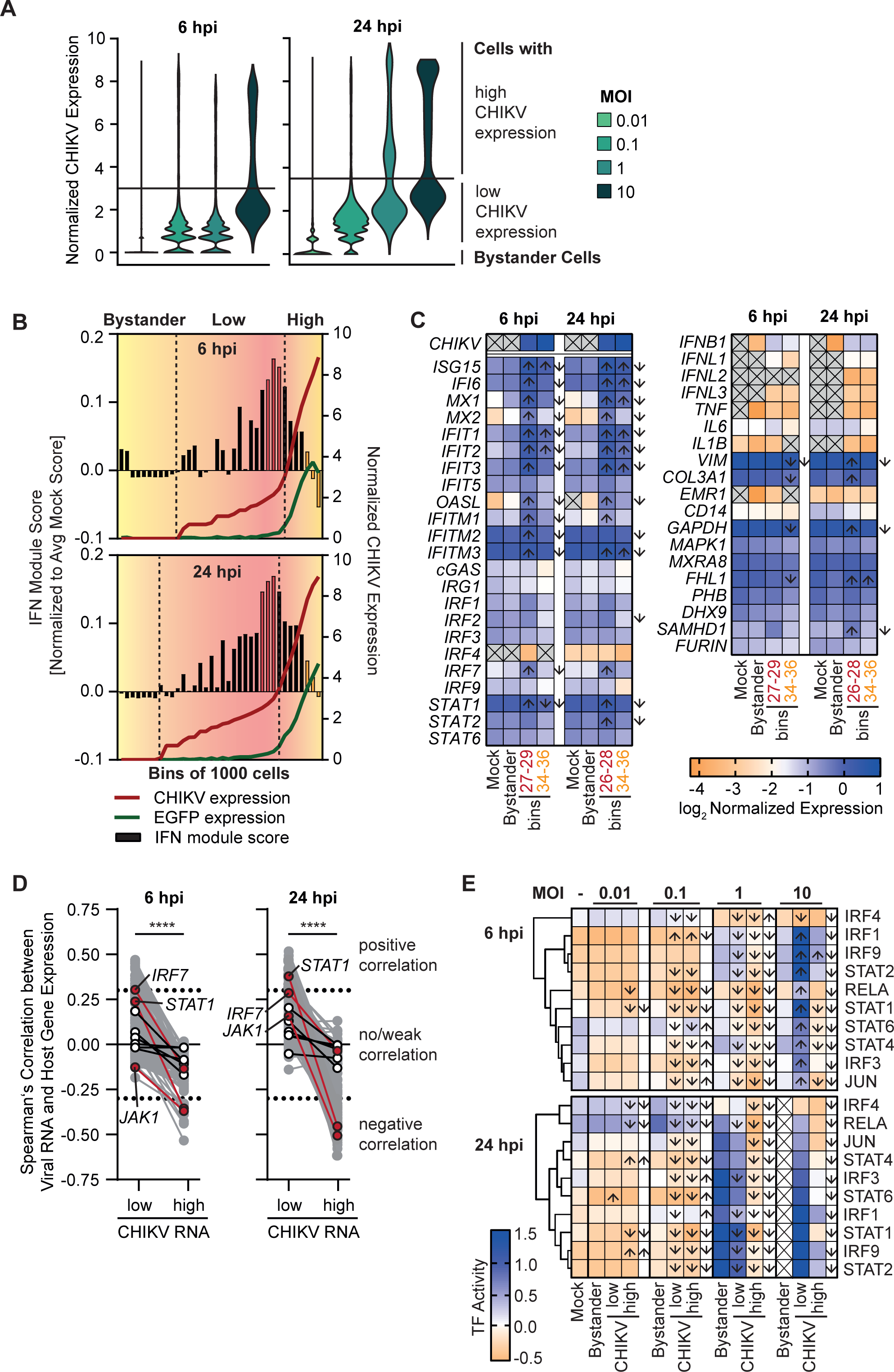
Transcriptomic differences between uninfected, bystander, and lowly or highly CHIKV infected OASF. (**A**) Visualization of the viral RNA content of infected OASF from Fig. 5 at six and 24 hours post-infection. Line indicates the cutoff dividing cells displaying low and high content of viral RNA. Bystander cells were defined as cells with no detectable viral RNA. (**B**) Infected OASF were sorted into digital bins of 1000 cells displaying a gradual increase of the amount of viral reads per cell. Viral reads and the IMS at six and 24 hours post-infection are plotted. Colored bins indicate selected representative cells for low and high content of viral RNA. (**C**) Expression of selected genes within mock, bystander, representative low cell bins (bin 26-29) and high cell bins (bin 34-36) defined in A and B at six and 24 hours post-infection. Arrows indicate a statistically significant (p<0.05, fold change >1.5) up- or downregulation (depending on the arrow direction) in low or high CHIKV bins versus bystander (inside the boxes) or in high CHIKV bins versus low (next to the boxes). (**D**) Correlation of CHIKV RNA expression with expression of IFN signaling genes in high and low CHIKV RNA groups calculated by non-parametric Spearman’s test. Transcription factors are plotted in white, with selected genes in red. (**E**) Activity of transcription factor regulons within groups defined in A at six and 24 hours post-infection. Arrows indicate a significant up- or downregulation between bystander and low or high CHIKV groups (inside the boxes) or between low and high CHIKV groups (next to the boxes).

To improve resolution, we calculated the average CHIKV and EGFP RNA expression and the average IMS in bins of 1000 cells for a total of 36 bins, sorted by their expression level of CHIKV RNA. At both time points, while the first 7-10 bins represented cells expressing no or virtually no CHIKV RNA, the following 18-21 bins represented cells displaying (according to the cut-off defined in Fig. 6A) low, but gradually increasing levels of CHIKV RNA, and largely undetectable EGFP RNA. We considered the latter cells to represent abortively infected cells due to their lack of subgenomic transcripts. The last eight bins displayed cells with overall high, starkly increasing levels of CHIKV RNA and with significant levels of EGFP mRNA. We hypothesize that these cells represent productively infected cells. Strikingly, in abortively infected cells, IMS values increased proportionally to the abundance of viral RNA per cell, whereas in productively infected cells, an inverse proportionality was observed (Fig. 6B). This dataset suggests that expression of IFN signaling genes is upregulated in cells harboring low-to-intermediate levels of viral RNA, which, however, do not or have not yet progressed to a productive infection. In contrast, cells that exceed a certain threshold of viral RNA show a prevention or downregulation of the expression of IFN signaling genes. The analysis of expression of selected genes confirmed this observation. Expression of individual ISGs, including including *ISG15*, *IFI6*, *MX1*, *OASL*, *IFITs*, and *IFITMs,* and transcription factors, including *STAT1* and *IRF7*, was low in mock-infected and bystander cells, and more pronounced in a representative low cells (six hours post-infection: bin 27-29, 24 hours post-infection: bin 26-28) than in high cells (bins 34-36) (Fig. 6C). To identify further putative targets of viral antagonism, we correlated the expression of all 203 genes of the IMS to the viral RNA expression in infected cells at 24 hours post-infection. We identified 13 genes displaying a significant positive correlation (r > 0.3) in the low CHIKV group, and a significant negative correlation (r < -0.3) in the high CHIKV group (*IFITM3, IFIT3, OAS1, XAF1, GBP1, EIF4A1, EIF2AK2, STAT1, GBP3, UBC, PSMB8, UBA52*). Strikingly, the only transcription factor present in both groups, *STAT1*, was also negatively correlated at six hours post-infection in the high viral RNA group. We additionally identified transcription factors *JAK1* and *IRF7* to switch from weak correlation in the low CHIKV group to a negative correlation in the high CHIKV group at 24 hours post-infection (Fig. 6D). We confirmed this finding using a transcription factor activity score analysis using the DoRotheEa database, which scores cells based on the activity of transcription factors inferred from the expression of the associated target genes in regulons, and found the regulon of *STAT1* to be strongly induced in bystander and low CHIKV groups at high MOIs, yet highly susceptible to viral antagonism in the high CHIKV group. This was also true for multiple other transcription factors – IRFs and STATs as well as NFκB and c-JUN - at higher MOIs, indicating a strong and sensitive induction that is counteracted in highly infected cells (Fig. 6E). Taken together, we demonstrate that the interaction between the virus and the host cell can be defined more precisely at single-cell resolution than by analyzing bulk data, and that the activation of innate immune responses can be well defined and correlated to the amount of viral RNA in the cell. Furthermore, viral antagonism may be masked by strong immune responses in cells infected at low levels, making it difficult to analyze using conventional RNA-seq.

## Discussion

Considering the CHIKV-induced arthritis, it is likely that cells of the synovium are directly implicated in the pathophysiology of CHIKV infection. Cells of the synovial tissue and synovial fluid contain CHIKV RNA and protein upon CHIKV infection *in vivo* in humans [9], experimentally infected macaques [48], and mice [49]. The main cell types composing the synovium are macrophages and fibroblasts. The latter have been identified to be susceptible to CHIKV infection *ex vivo* [12,50,51]. However, the corresponding basal innate immune state of primary synovial fibroblasts and their ability to exert IFN-mediated antiviral restriction is unknown. Here, we establish that the widely available OASF and less available HSF share susceptibility and permissiveness to CHIKV infection, and describe their basal and infection-induced transcriptional programs. These findings are in line with reports on overall transcriptional similarity of the two cell types, except in some signaling pathways unrelated to immunity [52]. CHIKV infection provoked a striking cellular response that involves upregulation of multiple ISGs, many of them exerting antiviral activity. Although we did not define the PAMP(s) that trigger responses in synovial macrophages, infection by alphaviruses typically raises RIG-I-mediated responses through exposure of dsRNA intermediates and provokes mitochondrial DNA leakage that is sensed via cGAS/STING [44,53,54]. Indeed, experimental ligands of both sensors were highly reactive in OASF, as was IFN-α treatment. Surprisingly, also IFN-λ pre-treatment translated into an antiviral state, indicating that synovial fibroblasts may represent an exception to the notion of otherwise IFN-λ-nonresponsive fibroblasts [55]. Finally, CHIKV infection of synovial fibroblasts was sensitive to IFN-α applied after inoculation with virus. These findings appear to contrast with potent virus-mediated antagonism of IFN in U2OS and HFF-1 cell lines, which has been suggested to involve counteraction of nuclear translocation of STAT1 [17, 18]. CHIKV was unable to suppress ISG expression upon exogenous IFN treatment in any cell type, indicating that the proposed antagonistic functions may not be strong enough to be detectable at the bulk level. Also, unaltered levels of expression of housekeeping genes and genes encoding fibroblast markers in primary synovial fibroblasts did not generate evidence for a general virus-mediated host transcriptional shut-off that has been reported for several cell lines [19, 20]. Overall, synovial fibroblasts appear to respond differently to CHIKV infection as commonly used cell lines. The underlying reason for this difference is unknown, but may involve a different intracellular milieu that is hyper-responsive to CHIKV infection.

While our single-cell RNA-seq approach is dependent on 3’ end capture and does not allow for the discrimination between full-length and partial viral RNA, we found an excess of subgenomic RNA in infected cells. We found a similar the ratio of subgenomic to genomic RNA as measured in Sindbis virus infected cells [56]. The enhanced replication of the subgenomic RNA, which is mediated by the four cleaved nonstructural proteins forming a replication complex, ensures the rapid production of viral structural proteins and the formation of new virions [57, 58]. While packaging of subgenomic RNA into virions has been described so far for one alphavirus, Aura virus [59], CHIKV holds a packaging signal in the nsP2 region of its genome, selecting only for full genomic RNA to be packaged into virions [60]. Therefore, we assume that the different abundance is based on de *novo* produced subgenomic RNA rather than on incoming viral RNA.

Through correlating cellular gene expression with CHIKV RNA abundance in individual infected cells, it appears that a certain threshold of viral RNA is required to initiate viral RNA sensing and eventually trigger ISG expression. However, expression of most ISGs is negatively regulated in the presence of a high viral RNA burden per cell. This is fully consistent with the idea that productive infection involves the synthesis of viral antagonists that hamper the induction and/or evade the function of ISGs, resulting in efficient virus propagation. Along these lines, West Nile virus infection also results in lowered ISG expression levels in cells harboring high viral RNA quantities [61]. *In vivo,* actively SARS-CoV-2 infected monocytes of COVID-19 patients expressed lower levels of ISGs than non-infected bystander [62]. Monocytes of Ebola-infected rhesus monkeys display similar dynamics, with an additional downregulation of *STAT1* mRNA in infected cells [63]. On the contrary, cells that undergo abortive infection, or alternatively haveńt yet reached sufficient levels of virus replication, fail to mount a strong antiviral profile. HIV-1 infection of lymphoid resting T-cells, that provide a sub-optimal environment for HIV-1 infection, has been reported to result in the accumulation of abortive viral cDNA products that are sensed by IFI16 in an inflammasome/pyroptosis-dependent manner [64]. HSV-1 infection results in antiviral signaling specifically in cells in which replication is stalled and that display relatively low levels of viral gene expression [65]. Owing to genetic recombination and low fidelity of the alphaviral RNA-dependent polymerase, defective alphaviral genomes (DVGs) and defective alphaviral particles arise during virus replication, but are themselves replication-incompetent [66, 67]. Of note, our virus-inclusive sequencing approach does not have the power to distinguish between full-length viral genomes and defective or otherwise dead-end genomes. It will be interesting to test the contribution of the latter to triggering the strong cell-intrinsic innate recognition that we linked here to high intracellular viral RNA quantities in general. Strikingly, we find indication that the expression of some genes, such as proinflammatory transcription factors, may be actively targeted by CHIKV.

Finally, the interplay of tissue-resident, synovial macrophages and fibroblasts likely additionally modulates CHIKV infection and cellular responses. Macrophages have been found to productively infect primary human macrophages [68] and to harbor persistent viral RNA in a nonhuman primate infection model [48]. Furthermore, human synovial fibroblasts secrete cytokines such as IL-6, IL1B, and RANTES stimulating monocyte migration upon CHIKV infection, and drive them towards an osteoclast-like phenotype [69, 70]. Interestingly, we find a similar phenotype in infected fibroblasts with upregulation and/or secretion of IL-6 and RANTES, but not matrix-metalloproteases (MMPs), as described before [69]. This suggests an indirect role of synovial fibroblasts in the induction of arthralgia upon infection, however, a paracrine stimulation of MMP expression by infiltrating immune cells can not be excluded in this model. Bystander cells, defined here as cells from infected cultures without detectable viral RNA, may be strongly impacted through paracrine signaling by infected cells [71]. Interestingly, at 24 hours post-infection we do not observe an extensive activation of bystander cells in cultures infected with a low MOI, despite an established infection and a number of *IFNB*-expressing cells. On the other hand, at six hours post-infection in cultures infected with an MOI 10, a condition in which we expect that almost all cells have made contact with virus particles, we observe a strong activation of the RNA-negative cells. This indicates that either a rapid release of cytokines and interferons only in highly infected cultures, or an interferon-independent sensing of viral PAMPs leads to abortive infection.

It is tempting to speculate that the synovial fibroblast-specific hyperreactivity is linked to the long-term arthralgia observed *in vivo* in chronic CHIKV patients, and that pharmacological interference with hyperinflammation represents a feasible intervention approach towards the alleviation of long-term arthralgia. In rheumatoid arthritis, hyperactivated synovial fibroblasts invade the joint matrix, destroying/disrupting the cartilage and causing long-term inflammation [28, 72]. This and the subsequent attraction of immune cells, including monocyte-derived macrophages to the damaged sites, may represent important events in the progression to long-term morbidity [73]. Indeed, data obtained in recent clinical studies suggests that treatment of chikungunya-induced arthritis with the immunosuppressant methotrexate may be a beneficial strategy [74, 75]. The data presented here does not fully support the hypothesis that infected synovial fibroblasts display a phenotype similar to fibroblasts in rheumatoid arthritis, but key features such as the IL-1β-mediated IL-6 release, the aggressive proinflammatory gene expression in productively infected cells, and the strong expression of important cofactors make them likely to contribute to viral replication and disease progression *in vivo*.

## Author contributions

FP and CG conceptualized the study, designed experiments, and interpreted the data. FP and CG wrote the manuscript with input from all co-authors. FP performed all experiments. EN extracted the fibroblasts and performed the first culturing passages. DP, RJPB, and EW performed RNA-seq alignments and bioinformatic analysis. DP wrote major parts of the code used for single-cell RNA-seq analysis and performed the transcription factor analysis. CG, ML, and TP supervised the study and acquired funding. All authors critically discussed the findings and approved of the final version of the manuscript.

## Acknowledgements

We thank the sequencing core of the Helmholtz Centre for Infection Research (HZI) in Braunschweig and the Genomics platform of the Berlin Institute of Health (BIH) for preparation of the Illumina sequencing libraries and the next generation sequencing. Additionally, we thank the sequencing facility of the Max Delbrück Center for Molecular Medicine for the next generation sequencing and bioinformatic support. We thank M. Diamond for providing the MXRA8-Fc proteins. We thank Theresia Stradal, Jens Bohne, and Sandra Pellegrini for providing the U2OS cells, HEK293T cells, and HL116 cells, respectively. We thank Thomas Pietschmann, Institute for Experimental Virology, TWINCORE, and Christian Drosten for constant support. This work was supported by funding from Deutsche Forschungsgemeinschaft (DFG) to CG (GO2153/3-1; GO2153/6-1), by the Impulse and Networking Fund of the Helmholtz Association through the HGF-EU partnering grant PIE-008 to CG, and by funding from the Helmholtz Center for Infection Research (HZI) and Berlin Institute of Health (BIH) to CG.

---

**Supplemental Figure 1.**
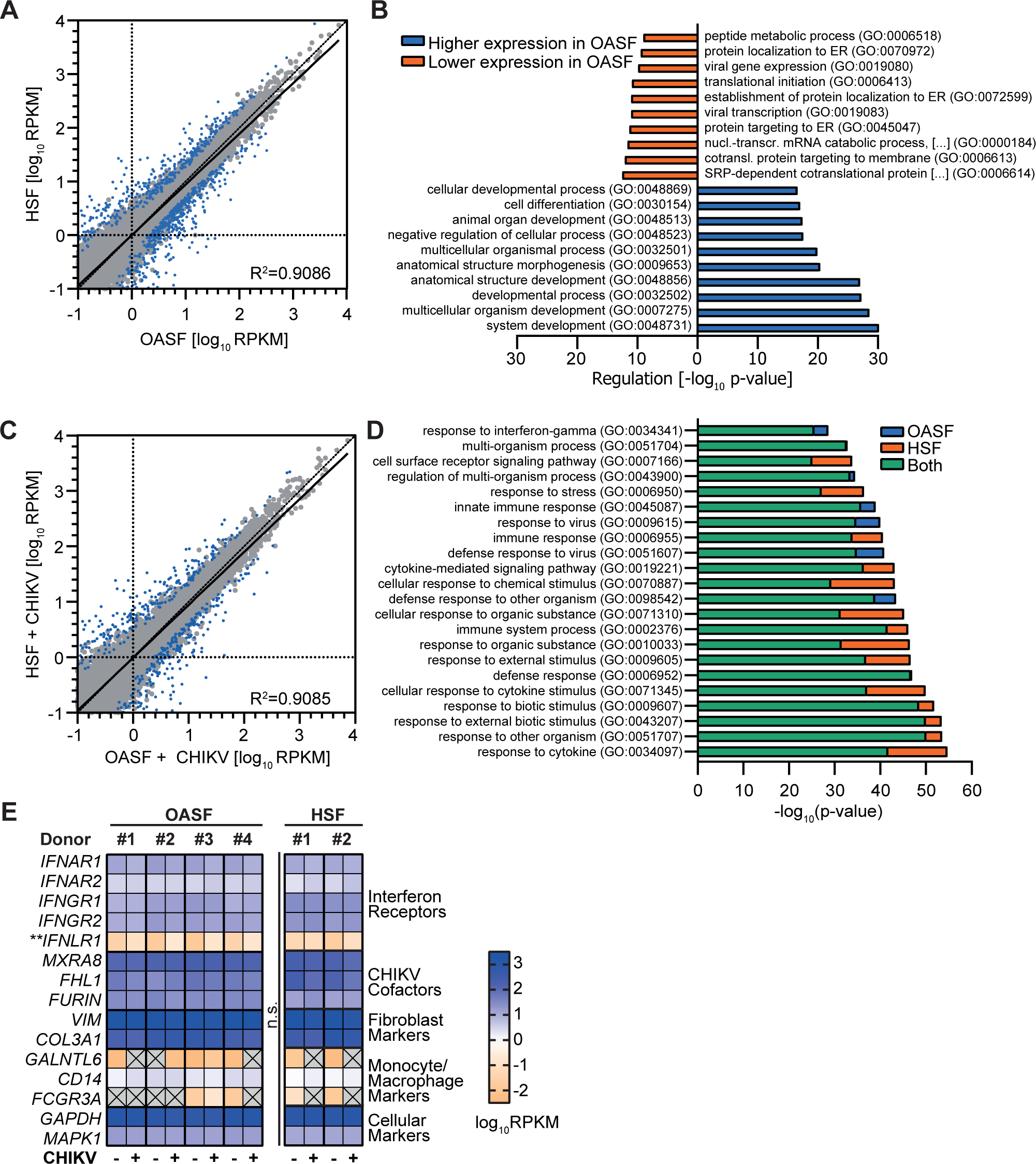
HSF and OASF share a similar basal and CHIKV infection-induced transcriptome. (**A**) Visualization of global transcriptional differences between OASF and HSF under regular culturing conditions. Average RPKM (log10) values for all detected transcripts from OASF are plotted on the x-axis, with corresponding values from HSF plotted on the y-axis. R^2^ value and regression line for comparison are inset. (**B**) Gene ontology analysis of differentially expressed genes in OASF compared to HSF. (**C**) Visualization of global transcriptomic differences between CHIKV-infected OASF and HSF as described in A. (**D**) Gene ontology analysis of the top significantly upregulated pathways in OASF, HSF, and shared by both in response to CHIKV infection. (**E**) Heatmaps of selected gene expression profiles of IFN receptors, CHIKV host cofactors, and celltype markers.

**Supplemental Figure 2.**
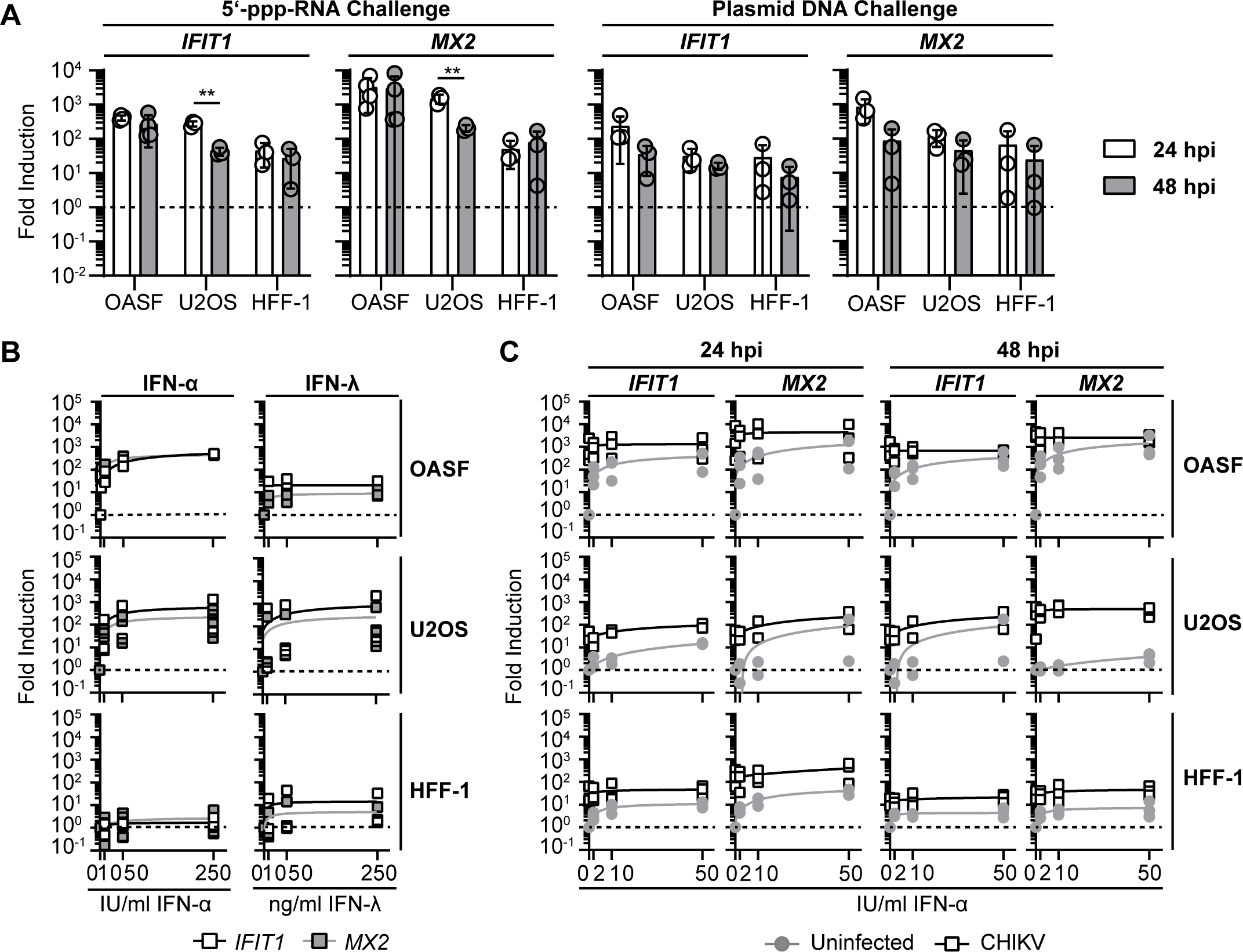
OASF, U2OS, and HFF-1 respond with a differently strong upregulation of ISGs to IFN treatment and CHIKV infection despite similar responsiveness to PAMPs. (**A**) Indicated cell cultures were transfected with 5’-triphosphate dsRNA (5-ppp-RNA, left) or plasmid DNA (right) and analyzed for the expression of *IFIT1* (left) and *MX2* (right) mRNA at 24 and 48 hours post transfection (n = 3-4). (**B**) OASF, U2OS, and HFF-1 cells were analyzed for the expression of *IFIT1* and *MX2* by quantitative RT-PCR after 48 h treatment with the indicated amounts of IFN-α or –λ. (**C**) OASF, U2OS, and HFF-1 cells were infected with 5’-EGFP CHIKV (MOI 10) and indicated amounts of IFN-α were added four hours post-infection. At 24 and 48 hours post-infection, OASF were analyzed for the expression of *IFIT1* and *MX2* mRNA by quantitative RT-PCR (n = 3).

**Supplemental Figure 3.**
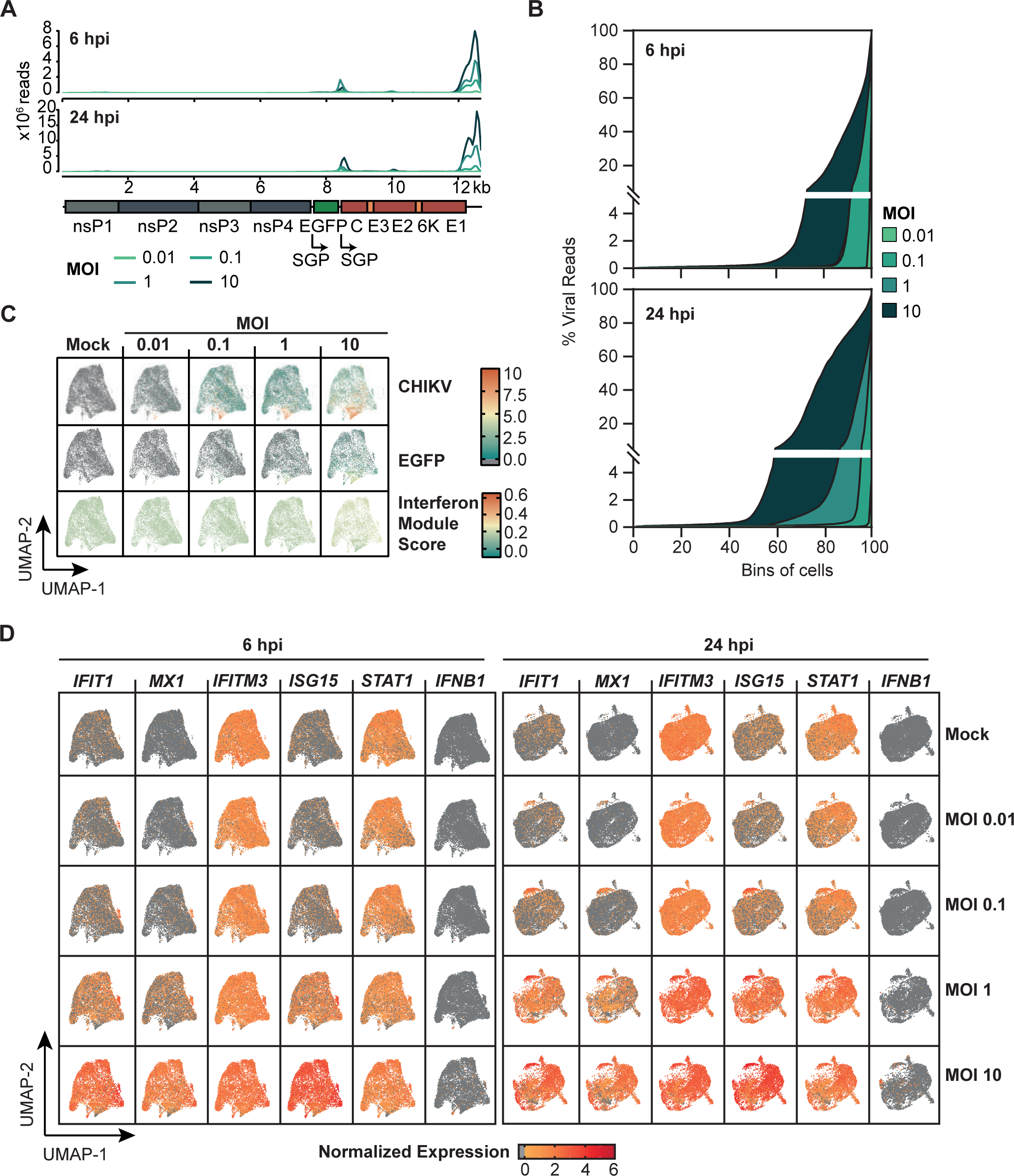
Cofactor expression in OASF and mild IFN signaling gene expression in infected OASF. (**A**) NGS reads after 3’ mRNA capture attributed to each individual position in the CHIKV genome plotted for cells infected with CHIKV at six and 24 hours post-infection. SGP = subgenomic promotor (**B**) Infected OASF were sorted into 100 digital bins per infection condition (MOI) displaying a gradual increase of the amount of viral reads per cell. Average proportion of reads per cell attributed to CHIKV at six and 24 hours post infection are plotted. (**C**) UMAP projections were generated for the six hours post infection time point. Shown is the abundance of CHIKV 3’ end reads and of IFN signaling genes according to IMS. (**D**) Expression of IFN-stimulated genes in infected OASF at six and 24 hours post-infection. UMAP visualization shows infected cells split by the MOI and mock-infected cells separately.

**Supplemental Figure 4.**
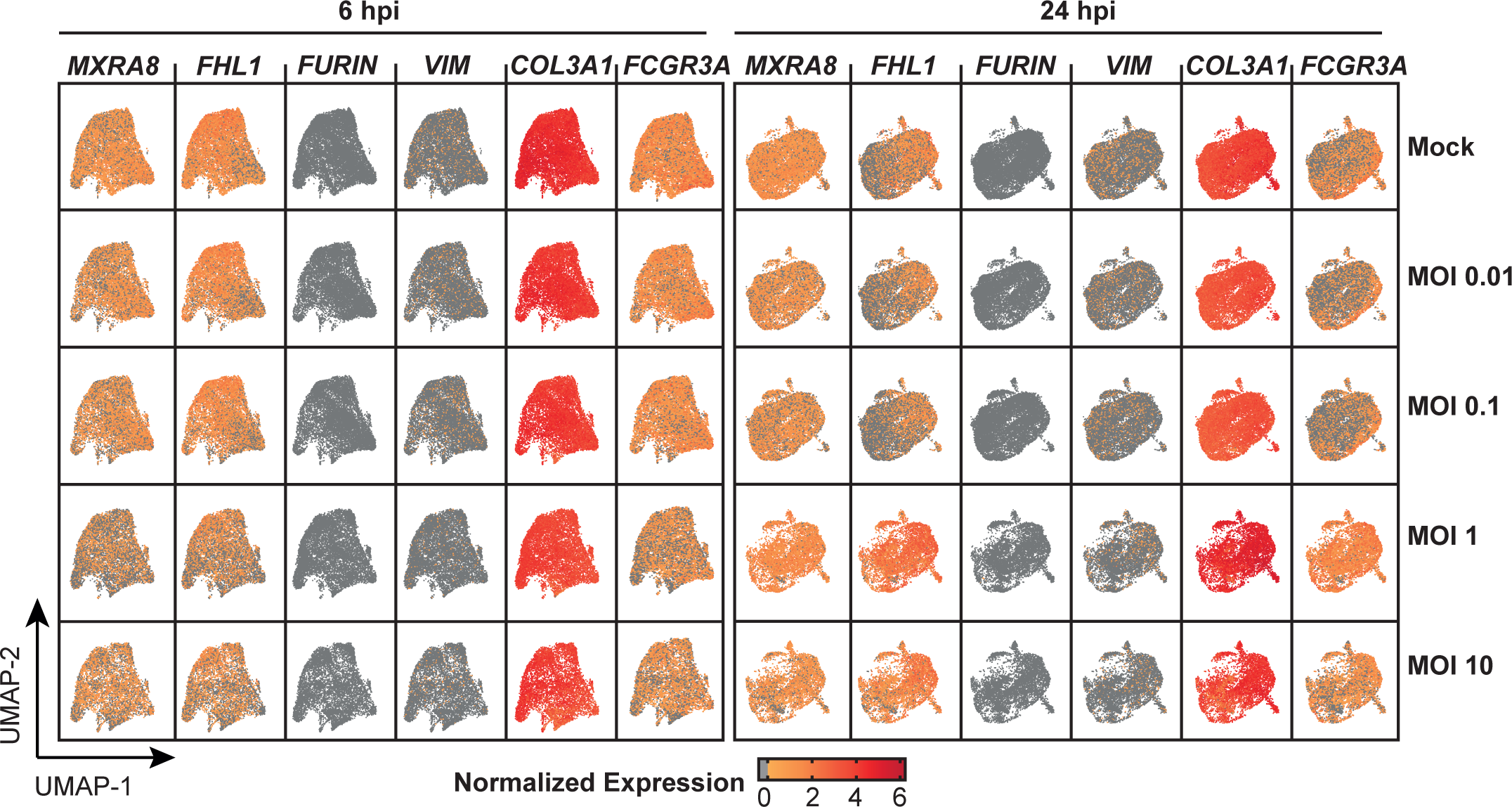
IFN-stimulated gene expression in infected OASF. Expression of alphavirus infection cofactors and fibroblast marker genes in infected OASF. UMAP visualization shows infected cells split by the MOI and mock-infected cells separately.

**Supplemental Figure 5.**
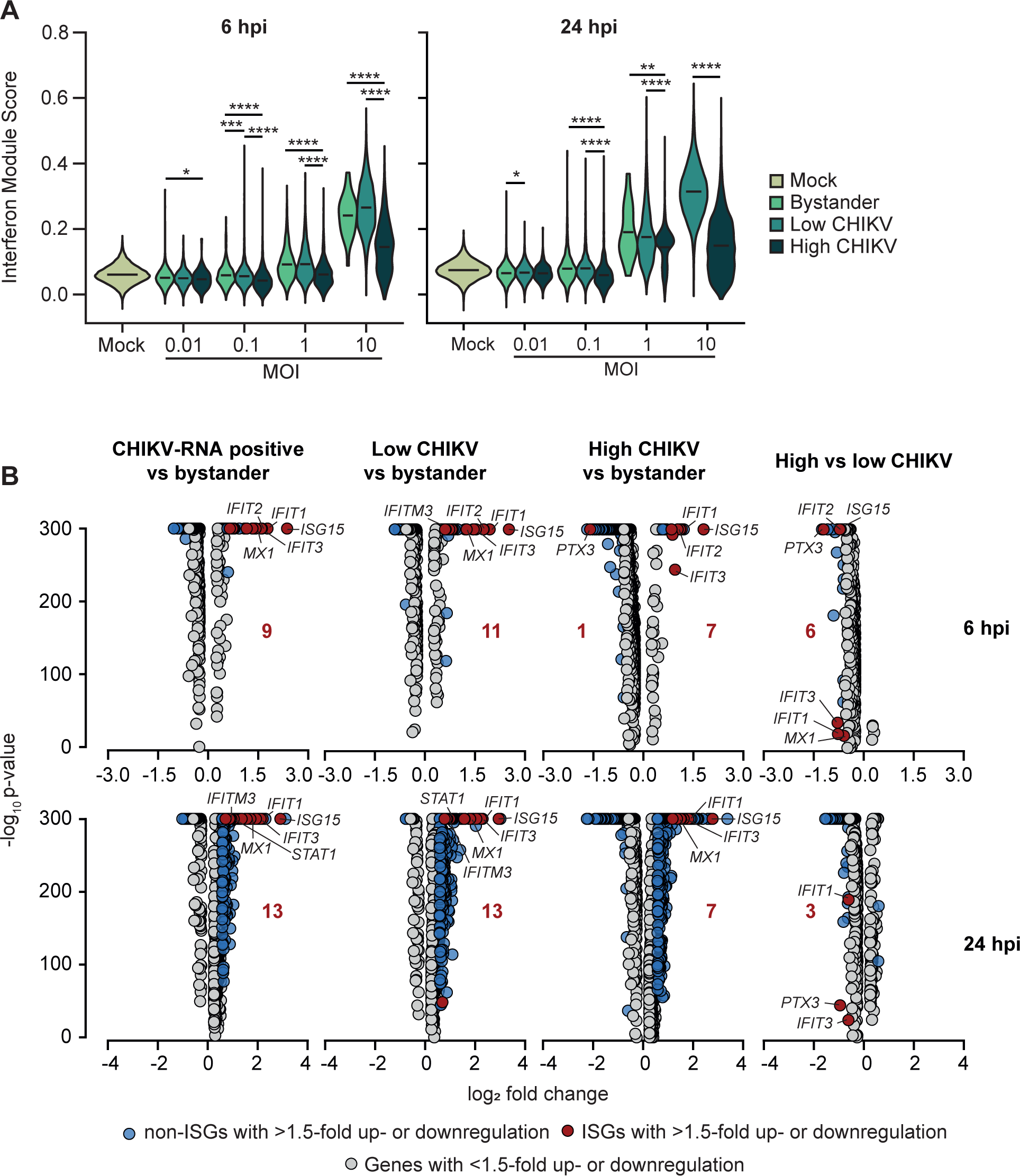
IFN signaling gene expression and transcriptional changes between infected subgroups of cells. (**A**) IMS in uninfected, bystander, CHIKV low and CHIKV high OASF at six and 24 hours post-infection. Statistical significance between groups was tested using a non-parametric KS-test. (**B**) Analysis of significantly up- and downregulated genes between indicated cell subgroups at six and 24 hours post-infection.

**Supplemental Movie 1. CHIKV spreads in OASF culture.**

OASF 5’-EGFP-CHIKV-infected OASF were monitored for EGFP expression by live-cell imaging. Scale bar = 100 μm.

**Supplemental Movie 2. Ruxolitinib treatment boosts CHIKV spread.**

OASF were pretreated with 10 μM Ruxolitinib for 16 hours, infected with 5’-EGFP-CHIKV in the presence of Ruxolitinib and monitored for EGFP expression by live-cell imaging. Scale bar = 100 μm.

## References

1. Matusali G, Colavita F, Bordi L, et al. Tropism of the Chikungunya Virus. Viruses. 2019 Feb 20;11(2).

2. Diagne CT, Bengue M, Choumet V, et al. Mayaro Virus Pathogenesis and Transmission Mechanisms. Pathogens. 2020 Sep 8;9(9).

3. Levi LI, Vignuzzi M. Arthritogenic Alphaviruses: A Worldwide Emerging Threat? Microorganisms. Vol. 72019.

4. Couderc T, Lecuit M. Chikungunya virus pathogenesis: From bedside to bench. Antiviral Res. 2015 Sep;121:120–31.

5. Paixao ES, Rodrigues LC, Costa M, et al. Chikungunya chronic disease: a systematic review and meta-analysis. Trans R Soc Trop Med Hyg. 2018 Jul 1;112(7):301–316.

6. Santiago FW, Halsey ES, Siles C, et al. Long-Term Arthralgia after Mayaro Virus Infection Correlates with Sustained Pro-inflammatory Cytokine Response. PLoS Negl Trop Dis. 2015;9(10):e0004104.

7. Suhrbier A. Rheumatic manifestations of chikungunya: emerging concepts and interventions. Nat Rev Rheumatol. 2019 Oct;15(10):597–611.

8. Chow A, Her Z, Ong EK, et al. Persistent arthralgia induced by Chikungunya virus infection is associated with interleukin-6 and granulocyte macrophage colony-stimulating factor. The Journal of infectious diseases. 2011 Jan 15;203(2):149–57.

9. Hoarau JJ, Jaffar Bandjee MC, Krejbich Trotot P, et al. Persistent chronic inflammation and infection by Chikungunya arthritogenic alphavirus in spite of a robust host immune response. J Immunol. 2010 May 15;184(10):5914–27.

10. Young AR, Locke MC, Cook LE, et al. Dermal and muscle fibroblasts and skeletal myofibers survive chikungunya virus infection and harbor persistent RNA. PLoS Pathog. 2019 Aug;15(8):e1007993.

11. Chang AY, Martins KAO, Encinales L, et al. Chikungunya Arthritis Mechanisms in the Americas: A Cross-Sectional Analysis of Chikungunya Arthritis Patients Twenty-Two Months After Infection Demonstrating No Detectable Viral Persistence in Synovial Fluid. Arthritis Rheumatol. 2018 Apr;70(4):585–593.

12. Zhang R, Kim AS, Fox JM, et al. Mxra8 is a receptor for multiple arthritogenic alphaviruses. Nature. 2018 May;557(7706):570–574.

13. Meertens L, Hafirassou ML, Couderc T, et al. FHL1 is a major host factor for chikungunya virus infection. Nature. 2019 Sep 25.

14. Salvador B, Zhou Y, Michault A, et al. Characterization of Chikungunya pseudotyped viruses: Identification of refractory cell lines and demonstration of cellular tropism differences mediated by mutations in E1 glycoprotein. Virology. 2009 Oct 10;393(1):33–41.

15. Reynaud JM, Kim DY, Atasheva S, et al. IFIT1 Differentially Interferes with Translation and Replication of Alphavirus Genomes and Promotes Induction of Type I Interferon. PLoS Pathog. 2015 Apr;11(4):e1004863.

16. Poddar S, Hyde JL, Gorman MJ, et al. The Interferon-Stimulated Gene IFITM3 Restricts Infection and Pathogenesis of Arthritogenic and Encephalitic Alphaviruses. J Virol. 2016 Oct 1;90(19):8780–94.

17. Fros JJ, Liu WJ, Prow NA, et al. Chikungunya virus nonstructural protein 2 inhibits type I/II interferon-stimulated JAK-STAT signaling. J Virol. 2010 Oct;84(20):10877–87.

18. Goertz GP, McNally KL, Robertson SJ, et al. The Methyltransferase-Like Domain of Chikungunya Virus nsP2 Inhibits the Interferon Response by Promoting the Nuclear Export of STAT1. J Virol. 2018 Sep 1;92(17).

19. Akhrymuk I, Kulemzin SV, Frolova EI. Evasion of the innate immune response: the Old World alphavirus nsP2 protein induces rapid degradation of Rpb1, a catalytic subunit of RNA polymerase II. Journal of virology. 2012 Jul;86(13):7180–91.

20. Akhrymuk I, Lukash T, Frolov I, et al. Novel Mutations in nsP2 Abolish Chikungunya Virus-Induced Transcriptional Shutoff and Make the Virus Less Cytopathic without Affecting Its Replication Rates. Journal of virology. 2019 Feb 15;93(4).

21. Haese NN, Broeckel RM, Hawman DW, et al. Animal Models of Chikungunya Virus Infection and Disease. The Journal of infectious diseases. 2016 Dec 15;214(suppl 5):S482–s487.

22. Simarmata D, Ng DC, Kam YW, et al. Early clearance of Chikungunya virus in children is associated with a strong innate immune response. Scientific reports. 2016 May 16;6:26097.

23. Schilte C, Couderc T, Chretien F, et al. Type I IFN controls chikungunya virus via its action on nonhematopoietic cells. J Exp Med. 2010 Feb 15;207(2):429–42.

24. Soares-Schanoski A, Baptista Cruz N, de Castro-Jorge LA, et al. Systems analysis of subjects acutely infected with the Chikungunya virus. PLoS Pathog. 2019 Jun;15(6):e1007880.

25. Hussain KM, Lee RC, Ng MM, et al. Establishment of a Novel Primary Human Skeletal Myoblast Cellular Model for Chikungunya Virus Infection and Pathogenesis. Scientific reports. 2016 Feb 19;6:21406.

26. Sukkaew A, Thanagith M, Thongsakulprasert T, et al. Heterogeneity of clinical isolates of chikungunya virus and its impact on the responses of primary human fibroblast-like synoviocytes. J Gen Virol. 2018 Apr;99(4):525–535.

27. Bernard E, Hamel R, Neyret A, et al. Human keratinocytes restrict chikungunya virus replication at a post-fusion step. Virology. 2015 Feb;476:1–10.

28. Neumann E, Lefevre S, Zimmermann B, et al. Rheumatoid arthritis progression mediated by activated synovial fibroblasts. Trends Mol Med. 2010 Oct;16(10):458–68.

29. Lefevre S, Meier FM, Neumann E, et al. Role of synovial fibroblasts in rheumatoid arthritis. Curr Pharm Des. 2015;21(2):130–41.

30. Uzé G, Di Marco S, Mouchel-Vielh E, et al. Domains of interaction between alpha interferon and its receptor components. J Mol Biol. 1994 Oct 21;243(2):245–57.

31. Neumann E, Riepl B, Knedla A, et al. Cell culture and passaging alters gene expression pattern and proliferation rate in rheumatoid arthritis synovial fibroblasts. Arthritis research & therapy. 2010;12(3):R83.

32. Tsetsarkin K, Higgs S, McGee CE, et al. Infectious clones of Chikungunya virus (La Reunion isolate) for vector competence studies. Vector borne and zoonotic diseases (Larchmont, NY). 2006 Winter;6(4):325–37.

33. Li X, Zhang H, Zhang Y, et al. Development of a rapid antiviral screening assay based on eGFP reporter virus of Mayaro virus. Antiviral Res. 2019 May 29;168:82–90.

34. The Gene Ontology Resource: 20 years and still GOing strong. Nucleic Acids Res. 2019 Jan 8;47(D1):D330–d338.

35. Ashburner M, Ball CA, Blake JA, et al. Gene ontology: tool for the unification of biology. The Gene Ontology Consortium. Nat Genet. 2000 May;25(1):25–9.

36. Hao Y, Hao S, Andersen-Nissen E, et al. Integrated analysis of multimodal single-cell data. bioRxiv. 2020:2020.10.12.335331.

37. Holland CH, Tanevski J, Perales-Patón J, et al. Robustness and applicability of transcription factor and pathway analysis tools on single-cell RNA-seq data. Genome Biol. 2020 Feb 12;21(1):36.

38. Basore K, Department of Pathology & Immunology WUSoM, Saint Louis, MO 63110, USA, Kim AS, et al. Cryo-EM Structure of Chikungunya Virus in Complex with the Mxra8 Receptor. Cell. 2019;0(0).

39. Georganas C, Liu H, Perlman H, et al. Regulation of IL-6 and IL-8 expression in rheumatoid arthritis synovial fibroblasts: the dominant role for NF-kappa B but not C/EBP beta or c-Jun. J Immunol. 2000 Dec 15;165(12):7199–206.

40. Gitter BD, Labus JM, Lees SL, et al. Characteristics of human synovial fibroblast activation by IL-1 beta and TNF alpha. Immunology. 1989 Feb;66(2):196–200.

41. Kay J, Calabrese L. The role of interleukin-1 in the pathogenesis of rheumatoid arthritis. Rheumatology (Oxford). 2004 Jun;43 Suppl 3:iii2–iii9.

42. Selvarajah S, Sexton NR, Kahle KM, et al. A neutralizing monoclonal antibody targeting the acid-sensitive region in chikungunya virus E2 protects from disease. PLoS Negl Trop Dis. 2013;7(9):e2423.

43. Hornung V, Ellegast J, Kim S, et al. 5’-Triphosphate RNA is the ligand for RIG-I. Science. 2006 Nov 10;314(5801):994–7.

44. Sanchez David RY, Combredet C, Sismeiro O, et al. Comparative analysis of viral RNA signatures on different RIG-I-like receptors. eLife. 2016 Mar 24;5:e11275.

45. Lazear HM, Schoggins JW, Diamond MS. Shared and Distinct Functions of Type I and Type III Interferons. Immunity. 2019 Apr 16;50(4):907–923.

46. Schoggins JW, Wilson SJ, Panis M, et al. A diverse range of gene products are effectors of the type I interferon antiviral response. Nature. 2011 Apr 28;472(7344):481–5.

47. Zhou JH, Wang YN, Chang QY, et al. Type III Interferons in Viral Infection and Antiviral Immunity. Cell Physiol Biochem. 2018;51(1):173–185.

48. Labadie K, Larcher T, Joubert C, et al. Chikungunya disease in nonhuman primates involves long-term viral persistence in macrophages. J Clin Invest. 2010 Mar;120(3):894–906.

49. Couderc T, Chrétien F, Schilte C, et al. A mouse model for Chikungunya: young age and inefficient type-I interferon signaling are risk factors for severe disease. PLoS Pathog. 2008 Feb 8;4(2):e29.

50. Agrawal M, Pandey N, Rastogi M, et al. Chikungunya virus modulates the miRNA expression patterns in human synovial fibroblasts. J Med Virol. 2020 Feb;92(2):139–148.

51. Selvamani SP, Mishra R, Singh SK. Chikungunya virus exploits miR-146a to regulate NF-κB pathway in human synovial fibroblasts. PLoS One. 2014;9(8):e103624.

52. Del Rey MJ, Usategui A, Izquierdo E, et al. Transcriptome analysis reveals specific changes in osteoarthritis synovial fibroblasts. Annals of the rheumatic diseases. 2012 Feb;71(2):275–80.

53. Aguirre S, Luthra P, Sanchez-Aparicio MT, et al. Dengue virus NS2B protein targets cGAS for degradation and prevents mitochondrial DNA sensing during infection. Nat Microbiol. 2017 Mar 27;2:17037.

54. Schoggins JW, MacDuff DA, Imanaka N, et al. Pan-viral specificity of IFN-induced genes reveals new roles for cGAS in innate immunity. Nature. 2014 Jan 30;505(7485):691–5.

55. Sommereyns C, Paul S, Staeheli P, et al. IFN-lambda (IFN-lambda) is expressed in a tissue-dependent fashion and primarily acts on epithelial cells in vivo. PLoS Pathog. 2008 Mar 14;4(3):e1000017.

56. Lemm JA, Rümenapf T, Strauss EG, et al. Polypeptide requirements for assembly of functional Sindbis virus replication complexes: a model for the temporal regulation of minus- and plus-strand RNA synthesis. Embo j. 1994 Jun 15;13(12):2925–34.

57. Levis R, Schlesinger S, Huang HV. Promoter for Sindbis virus RNA-dependent subgenomic RNA transcription. Journal of virology. 1990 Apr;64(4):1726–33.

58. Rupp JC, Sokoloski KJ, Gebhart NN, et al. Alphavirus RNA synthesis and non-structural protein functions. J Gen Virol. 2015 Sep;96(9):2483–2500.

59. Rümenapf T, Strauss EG, Strauss JH. Subgenomic mRNA of Aura alphavirus is packaged into virions. Journal of virology. 1994 Jan;68(1):56–62.

60. Kim DY, Firth AE, Atasheva S, et al. Conservation of a packaging signal and the viral genome RNA packaging mechanism in alphavirus evolution. Journal of virology. 2011 Aug;85(16):8022–36.

61. O’Neal JT, Upadhyay AA, Wolabaugh A, et al. West Nile Virus-Inclusive Single-Cell RNA Sequencing Reveals Heterogeneity in the Type I Interferon Response within Single Cells. Journal of virology. 2019 Mar 15;93(6).

62. Bost P, Giladi A, Liu Y, et al. Host-Viral Infection Maps Reveal Signatures of Severe COVID-19 Patients. Cell. 2020 Jun 25;181(7):1475–1488.e12.

63. Kotliar D, Lin AE, Logue J, et al. Single-Cell Profiling of Ebola Virus Disease In Vivo Reveals Viral and Host Dynamics. Cell. 2020 Nov 2.

64. Monroe KM, Yang Z, Johnson JR, et al. IFI16 DNA sensor is required for death of lymphoid CD4 T cells abortively infected with HIV. Science (New York, NY). 2014 Jan 24;343(6169):428–32.

65. Drayman N, Patel P, Vistain L, et al. HSV-1 single-cell analysis reveals the activation of anti-viral and developmental programs in distinct sub-populations. eLife. 2019 May 15;8.

66. Poirier EZ, Mounce BC, Rozen-Gagnon K, et al. Low-Fidelity Polymerases of Alphaviruses Recombine at Higher Rates To Overproduce Defective Interfering Particles. Journal of virology. 2015 Dec 16;90(5):2446–54.

67. Levi LI, Rezelj VV, Henrion-Lacritick A, et al. Defective viral genomes from chikungunya virus are broad-spectrum antivirals and prevent virus dissemination in mosquitoes. PLoS pathogens. 2021 Feb;17(2):e1009110.

68. Sourisseau M, Schilte C, Casartelli N, et al. Characterization of reemerging chikungunya virus. PLoS pathogens. 2007 Jun;3(6):e89.

69. Phuklia W, Kasisith J, Modhiran N, et al. Osteoclastogenesis induced by CHIKV-infected fibroblast-like synoviocytes: a possible interplay between synoviocytes and monocytes/macrophages in CHIKV-induced arthralgia/arthritis. Virus research. 2013 Nov 6;177(2):179–88.

70. Ng LF, Chow A, Sun YJ, et al. IL-1beta, IL-6, and RANTES as biomarkers of Chikungunya severity. PloS one. 2009;4(1):e4261.

71. Cruz MA, Parks GD. La Crosse Virus Infection of Human Keratinocytes Leads to Interferon-Dependent Apoptosis of Bystander Non-Infected Cells In Vitro. Viruses. 2020 Feb 25;12(3).

72. Hillen J, Geyer C, Heitzmann M, et al. Structural cartilage damage attracts circulating rheumatoid arthritis synovial fibroblasts into affected joints. Arthritis research & therapy. 2017 Feb 28;19(1):40.

73. Falconer J, Murphy AN, Young SP, et al. Review: Synovial Cell Metabolism and Chronic Inflammation in Rheumatoid Arthritis. Arthritis Rheumatol. 2018 Jul;70(7):984–999.

74. Amaral JK, Sutaria R, Schoen RT. Treatment of Chronic Chikungunya Arthritis With Methotrexate: A Systematic Review. Arthritis Care Res (Hoboken). 2018 Oct;70(10):1501–1508.

75. Ganu MA, Ganu AS. Post-chikungunya chronic arthritis--our experience with DMARDs over two year follow up. J Assoc Physicians India. 2011 Feb;59:83–6.

